# Multiomics-driven discovery of predictive biomarkers and strategies to overcome resistance to SFK-YAP inhibition in cholangiocarcinoma

**DOI:** 10.64898/2026.01.21.699926

**Authors:** Hendrien Kuipers, Danielle M. Carlson, Erik Jessen, Aushinie M. Abeynayake, Jack W. Sample, Dong-Gi Mun, Jennifer L. Tomlinson, Amro M. Abdelrahman, Nathan W. Werneburg, Binbin Li, Enis H. Ozmert, Caitlin B. Conboy, Mitesh J. Borad, Mark J. Truty, Akhilesh Pandey, Sumera I. Ilyas, Gregory J. Gores, Rory L. Smoot

## Abstract

The limited efficacy of current therapies against cholangiocarcinoma (CCA) necessitates the development of novel treatment strategies. Src family kinases (SFKs) contribute significantly to tumor progression and resistance in CCA. Therefore, we investigated the novel, first-in-class SFK ‘OFF’ inhibitor NXP900 in diverse preclinical CCA models, including those with acquired resistance. This study evaluated the therapeutic effects of NXP900 and detailed adaptive molecular responses to SFK inhibitor therapy. We also aimed to identify biomarkers predictive of drug sensitivity using integrated multiomic profiling and develop strategies to overcome resistance. NXP900 inhibited YAP activity through direct inhibition of tyrosine phosphorylation and indirect activation of the Hippo pathway via LATS. These effects were associated with decreased tumor cell viability in CCA cell lines and several *in vivo* models. Notably, *IDH*-mutant patient-derived xenograft CCA models were particularly sensitive to NXP900. NXP900 also synergized with gemcitabine/cisplatin chemotherapy, enhancing antitumor efficacy in both *in vitro* and *in vivo* models. Multiomic analyses combining transcriptomics, global proteomics, and phosphoproteomics identified molecular features associated with primary response and acquired resistance. IL13RA-AKT signaling was upregulated in resistant models; NXP900 sensitivity could be restored with AKT or IL13RA2 inhibition. Together, these findings demonstrate the therapeutic potential of NXP900 as a novel YAP inhibitor in CCA and support further investigation in a clinical trial.

**Graphical Abstract:** 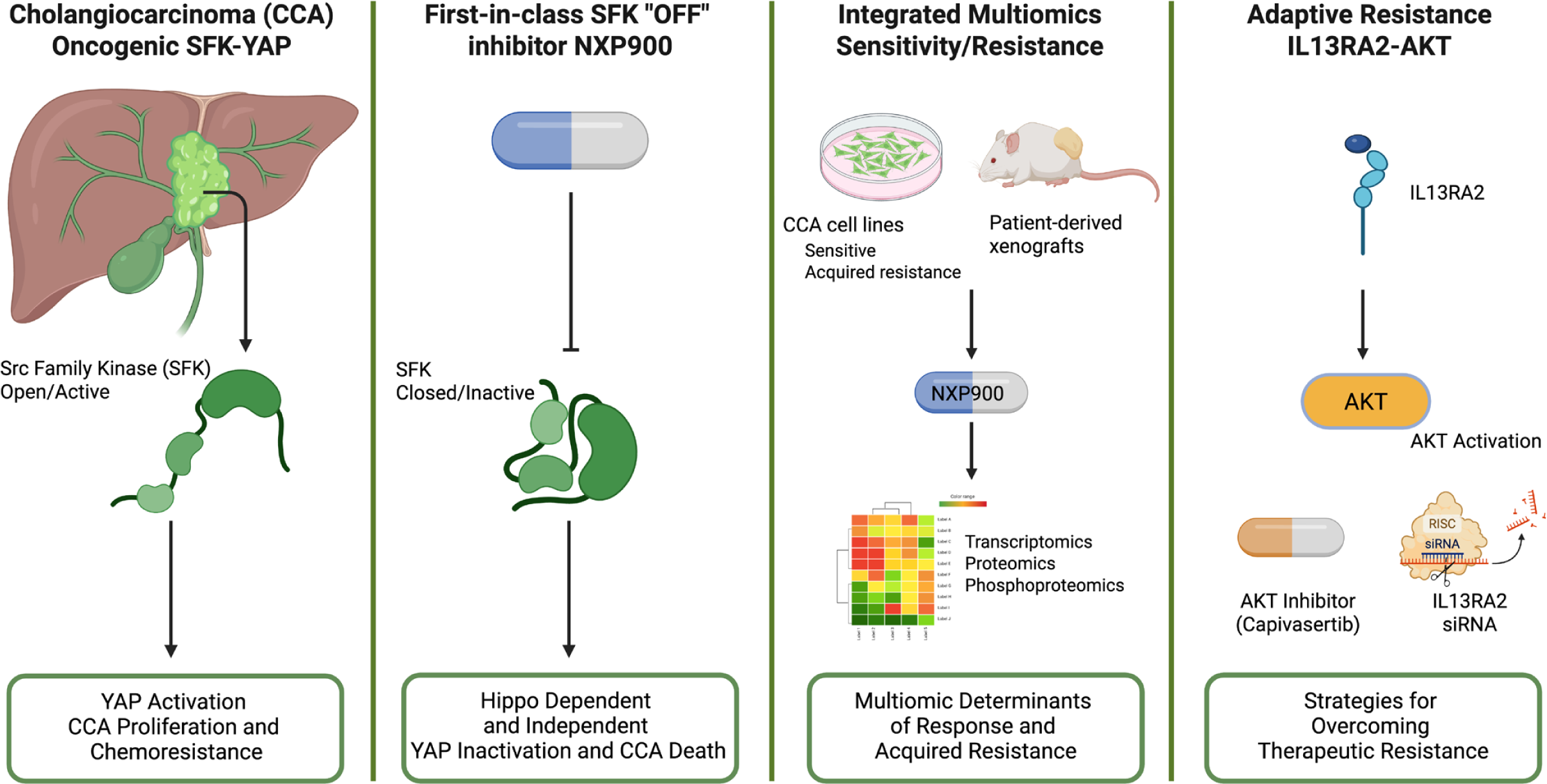

## INTRODUCTION

Cholangiocarcinoma (CCA) is a primary liver cancer characterized by differentiated features of the biliary epithelium. Based on anatomic origin, it can be classified as perihilar, intrahepatic, or distal CCA ^1^. The 5-year overall survival rate is only 9-11%, as the majority of patients are diagnosed with advanced or metastatic disease ^2^. Even following curative-intent surgery, recurrence rates are around 60%, suggesting the presence of micrometastatic disease in most patients and emphasizing the need for systemic therapeutic approaches ^3^. In the setting of advanced or unresectable disease, the current first-line therapy consists of gemcitabine, cisplatin, and either programmed death ligand 1 (PD-L1) (durvalumab) or programmed cell death protein 1 (PD-1) (pembrolizumab) blockade ^4–7^. Molecular characterization studies have identified targetable alterations such as fibroblast growth factor receptor (*FGFR)2* fusions or isocitrate dehydrogenase (*IDH)1* mutations in subsets of patients ^8–10^. However, even in patients with these molecular alterations, response rates and durability are limited ^11–13^. Therefore, the development of novel therapeutic strategies is urgently needed to improve outcomes for patients with CCA.

The Hippo pathway and its downstream effector Yes-associated protein (YAP) have been implicated in CCA pathogenesis ^14,15^. Approximately 85% of human CCA specimens demonstrate nuclear staining of YAP, indicative of co-transcriptional activity ^15–18^. The Hippo pathway plays a fundamental role in maintaining tissue homeostasis, regeneration and organ size through dynamic regulation. When active, it suppresses YAP activity, thereby restricting cell proliferation and promoting apoptosis; conversely, when downregulated, it promotes cell survival and proliferation ^19^. Due to its negative regulatory function, the Hippo pathway has been historically difficult to target and no approved targeted therapies exist ^19,20^. We have previously demonstrated a regulatory mechanism of YAP that is independent of the Hippo pathway and driven by an activating tyrosine phosphorylation and can be mediated by the SRC family kinases (SFKs) ^20–22^. SFKs are non-receptor protein tyrosine kinases which are overactivated in several cancer types and play a role in cancer cell survival, metastatic potential and resistance to therapies. Overexpression or activation of SFKs is associated with poor clinical prognosis in multiple cancers ^23–27^. We have previously shown that SFK inhibitors decrease YAP signaling and demonstrate preclinical efficacy in *in vitro* and *in vivo* CCA models ^21,22,28^. While previous studies have shown that SFK inhibition can be effective in *IDH-*mutant CCA models via dependency on SRC signaling, support from human clinical trials is lacking ^29–32^.

NXP900, formerly known as eCF506, is a first-in-class small molecule SFK ‘OFF’ inhibitor with the highest degree of inhibition to the SRC kinase in a kinome screening of 340 kinases ^33^. NXP900 has a novel mechanism of action which works as a switch-control that locks its target in the native closed and inactive (OFF) conformation, thus inhibiting both the catalytic activity and the scaffolding properties induced by oncogenic signaling ^33^. In this study, we investigated the effects of SFK inhibition utilizing NXP900 in CCA cell lines and patient-derived xenograft (PDX) models. We observed the anticipated suppression of YAP activation via direct inhibition of tyrosine phosphorylation. Furthermore, we noted activation of the Hippo pathway due to relief of SFK-mediated negative feedback on large tumor suppressor (LATS) activation. This dual inhibition of YAP, both directly and through the activation of the Hippo pathway, resulted in decreased YAP transcriptional activity, reduced CCA cell viability, and sensitization of CCA models to cytotoxic gemcitabine/cisplatin chemotherapy. Consistent with previous work, *IDH*-mutant CCA models were noted to be especially sensitive to NXP900 treatment. Multiomics approaches identified molecular features associated with primary response and non-response to NXP900 treatment and were subsequently utilized to query internal and external multiomic datasets to estimate the proportion of tumors likely to respond to single agent therapy. Finally, NXP900-resistant cell line models were generated and multiomic features associated with acquired resistance were defined and therapeutically targeted, resulting in restored sensitivity to NXP900 treatment.

## RESULTS

### NXP900 inhibits YAP activity and alters multiple signaling pathways in CCA cells

To evaluate the effect of NXP900 on CCA cell viability, we determined the half-maximal inhibitory concentration (IC50) in six human and murine CCA cell lines. All cell lines were sensitive to NXP900 in a concentration-dependent manner with IC50 values between 7 nM and 15 µM with cell viability approaching 0% at maximum drug concentrations (**Fig. 1A**). Mechanistically, cell death occurred via apoptosis, as demonstrated by increased caspase-3/7 activity (**Fig. 1B**).

**Figure 1.**
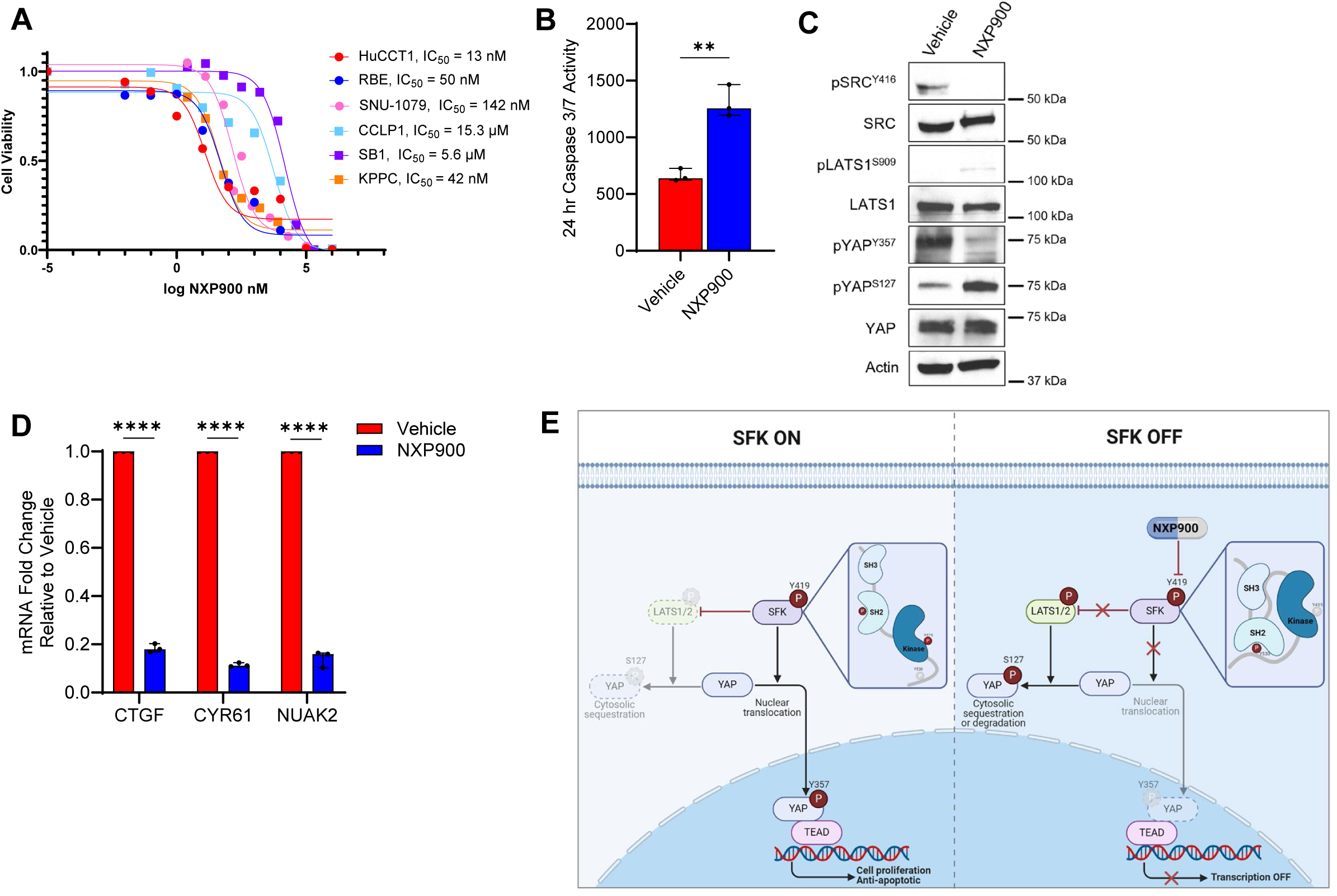
NXP900 induces apoptotic cell death and downregulates YAP signaling. (**A**) Viability dose-response curves and calculated IC50s of CCA cell lines treated with NXP900. (**B**) Caspase 3/7 activity in HuCCT1 cells after NXP900 (100 nM) treatment for 24 hours. (**C**) Cell lysates from the HuCCT1 cell line treated with NXP900 (1 µM for 6 hours) subjected to immunoblot for pSRCY416, total SRC, pLATS1S909, total LATS, pYAPY357, pYAPS127 and total YAP. Actin was used as a loading control. (**D**) mRNA expression in HuCCT1 cells after treatment with NXP900 (1 µM for 6 hours). (**E**) Proposed model of SFK/Hippo pathway interaction with the SFK inhibitor NXP900. Data are shown as median with 95% CI (*P < 0.05, **P < 0.01, ***P < 0.001, ****P < 0.0001). Statistical analysis was performed with 2-tailed Student t test.

Given that SRC inhibition reduces YAP Y357 phosphorylation, we examined the cellular effects of exposure to the SFK inhibitor NXP900 on YAP phosphorylation status and transcriptional activity. Accordingly, we observed decreased SRC Y416 and YAP Y357 phosphorylation levels after exposure to NXP900 (**Fig. 1C**). NXP900 relieved SRC-mediated inhibition of LATS leading to increased LATS S909 phosphorylation and subsequent increased YAP S127 phosphorylation, indicating that SRC modulates YAP through both Hippo-dependent and Hippo-independent mechanisms (**Fig. 1C**). NXP900 treatment was associated with a decrease in the abundance of the YAP target genes *CTGF, CYR61*, and *NUAK2 (***Fig. 1D**). These data suggest that NXP900 is therapeutically active in multiple CCA cell lines and effectively reduces YAP activation (**Fig. 1E**).

While we observed decreases in YAP activation in CCA models exposed to NXP900, we sought to further understand the landscape of molecular changes following SFK inhibition. We first exposed HuCCT1 cells to NXP900 (1 µM for 6 hours) and performed multiomic analysis integrating transcriptomics, global proteomics, and phosphoproteomics (**Suppl. Fig. 1**). Unsupervised clustering separated NXP900-treated and vehicle sample groups, highlighting induced molecular differences following treatment with NXP900 (**Fig. 2A**). RNA-Seq analysis demonstrated changes in multiple transcript levels in drug-treated cells compared to vehicle with several of the most differentially expressed genes being previously validated YAP target genes including *SERPINE1, ANKRD1, CTGF,* and *HBEGF* (an EGFR activator) (**Fig. 2B**, **Fig. 2C)** ^34–36^. SRC has been reported to mediate the bidirectional crosstalk of NGF–TrkA and EGFR pathways, creating a pro-survival signaling network ^37^. Disrupting SRC with NXP900 led to downregulation of *NGF*, NGF-responsive gene *EGR1*, and NGF/TrkA target gene *KLF2* (**Fig. 2B**, **Fig. 2C)** ^38^. The two most upregulated genes were the proapoptotic and anti-proliferative TP53 target genes, *TP53INP1* and *ARRDC3* (**Fig. 2B**, **Fig. 2C)** ^39–41^. Pathway analysis revealed decreased RAF/MAP kinase cascades and decreased RAF-independent MAPK1/3 activation, as well as enhanced cell cycle regulation upon treatment with NXP900 (**Fig. 2D**). Treatment with a structurally and mechanistically dissimilar pan-SFK inhibitor, dasatinib, was associated with similar transcriptomic changes (**Suppl. Fig. 2**). These findings were validated in a separate human CCA cell line, RBE. Similar to HuCCT1, RBE treated cells demonstrated downregulation of YAP target genes *SERPINE1*, *ANKRD1,* and *CTGF* following NXP900 exposure (**Fig. 2E**).

**Figure 2.**
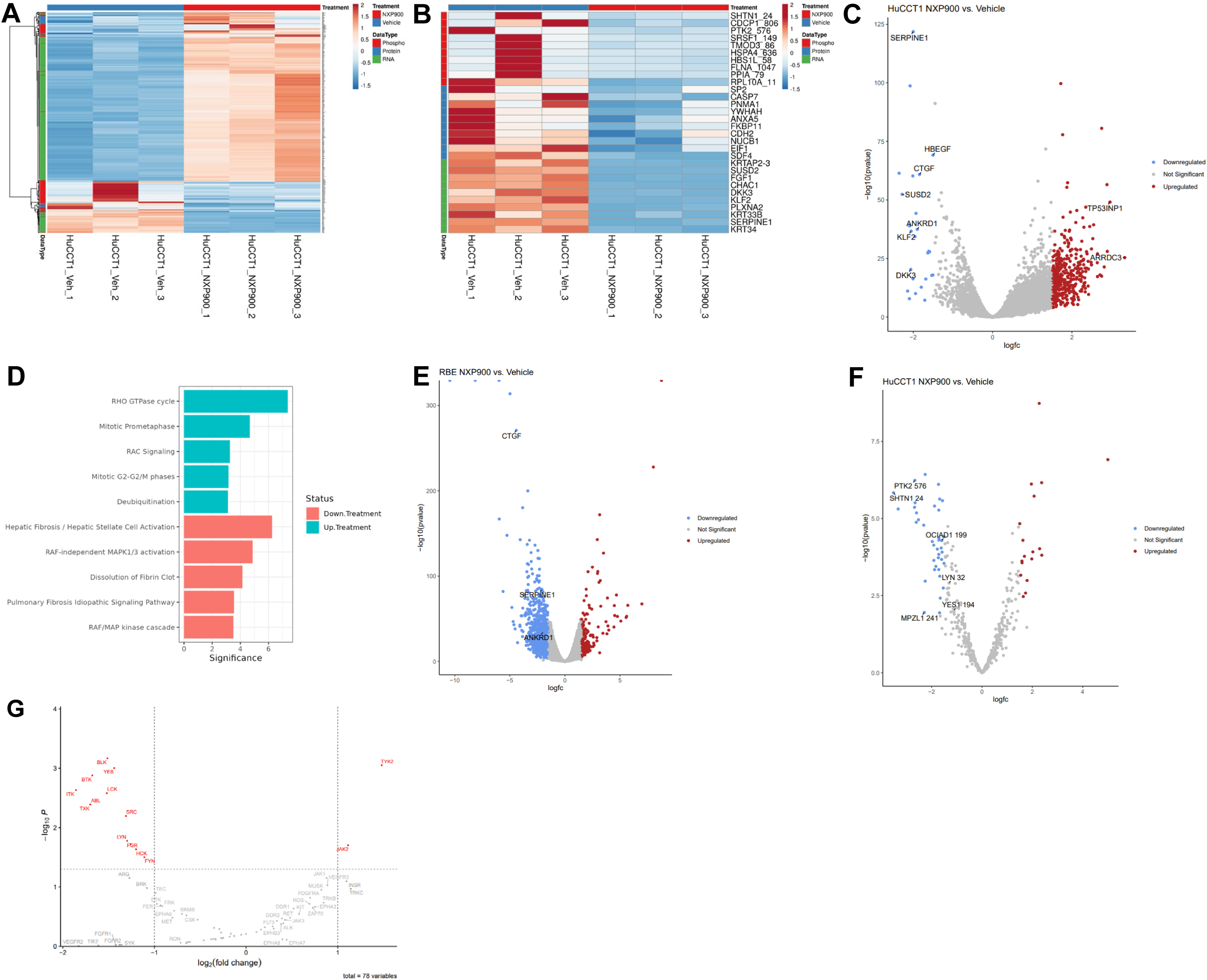
Multiomic in vitro characterization identifies treatment-related molecular alterations. Multiomic profiling of HuCCT1 cells (A-D, F-H) and RBE cells (E) treated with NXP900 (1 µM for 6 hours) compared to vehicle. (**A**) Broad characterization using RNA-Seq, global proteomics, and phosphoproteomics. (**B**) Top altered differentially expressed genes, proteins and phosphoproteins. (**C**) Volcano plot comparing RNA-Seq gene expression (3 technical replicates/group). Significantly differentially expressed genes (–1.5 < logFC > 1.5, FDR < 0.05) highlighted blue (downregulated) and red (upregulated) with top genes of interest identified on plot. (**D**) Canonical pathway analysis of the up-and downregulated pathways based on RNA-Seq results. Significance is the −log10(PValue) for the pathway. (**E**) Volcano plot comparing RNA-Seq gene expression in vehicle- and NXP900-treated RBE cells (3 technical replicates/group). Significantly differentially expressed genes (–1.5 < logFC > 1.5, FDR < 0.05) highlighted blue (downregulated) and red (upregulated) with top genes of interest identified on plot. (**F**) Volcano plot comparing phosphoproteomic (pTyr) abundance in vehicle- and NXP900-treated HuCCT1 cells (3 technical replicates/group). Significantly differentially expressed phosphosites (–1.5 < logFC > 1.5, FDR < 0.05) highlighted blue (downregulated) and red (upregulated) with top phosphoproteins of interest identified on plot. (**G**) Kinase enrichment analysis of Tyr phosphorylation data analysis using a kinase library (https://kinase-library.phosphosite.org/kinase-library/).

In parallel, we conducted phosphoproteomic (pTyr) and total proteomic analyses to assess changes after treatment with NXP900. **Fig. 2B** and **Fig. 2F** illustrate the phosphosites with increased and decreased abundance. Kinase enrichment analysis predicted decreased activity of SFK members (BLK, YES, SRC, LYN, FYN, HCK, LCK, FGR) as well as TEC kinases (BTK, ITK, TXK) following treatment with NXP900. ABL kinase activity was also decreased, likely as an off-target effect resulting from the short, high-dose regimen applied to minimize bypassing effects, which exceeded the IC90^33^. Activity of JAK2 and TYK2, key members of the JAK family, were predicted to be increased, suggesting an early compensatory response (**Fig. 2G**). There were no changes in global protein after NXP900 treatment for 6 hours.

These data demonstrate that NXP900 inhibits YAP transcriptional activity and favorably alters multiple signaling molecules, thereby suppressing oncogenic signaling.

### NXP900 is effective in vivo and can sensitize to first-line cytotoxic chemotherapy

Based on the effects seen in our cell line models, we next evaluated the efficacy and toxicity of NXP900 treatment *in vivo* using two syngeneic murine models of iCCA, KPPC (Kras^G12D^p53^L/L^) and SB1 (YAP^S127A^/myr-AKT) ^42,43^. Cells were orthotopically injected into the liver and after the engraftment period, tumor-bearing mice were treated with vehicle or NXP900 by oral gavage once daily (**Fig. 3A**). We observed a decrease in tumor size in NXP900-treated mice compared to vehicle in the KPPC model (**Fig. 3B**), but not in the SB1 model (**Fig. 3C)**. Of note, the SB1 cell line expresses an epitope-tagged YAP containing a S127A mutation as well as myr-AKT. The S127A mutation renders YAP insensitive to serine/threonine kinase phosphorylation, suggesting that the anti-tumor effects of SFK inhibition by NXP900 may be partially dependent on YAP S127 phosphorylation by serine/threonine kinases. Further, myr-AKT may alter resistance signaling in this cell line. Taken together these in vivo data mirror our *in vitro* findings and the high IC50 measured for the SB1 cell line (**Fig. 1A**).

**Figure 3.**
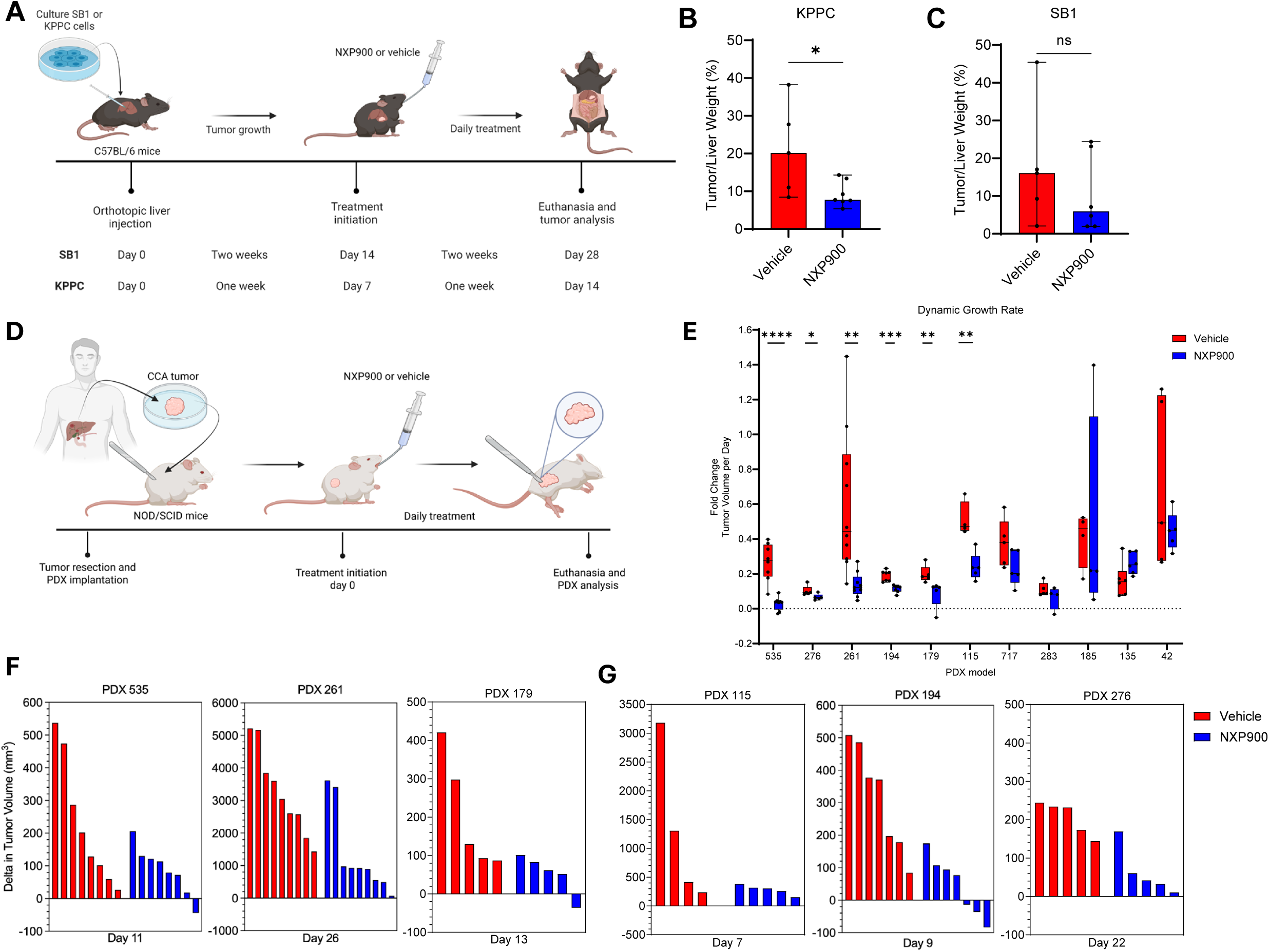
NXP900 is therapeutic in multiple in vivo models of CCA. (**A**) Schematic depicting orthotopic implantation model of SB1 or KPPC cells and subsequent tumor development and treatment with NXP900 or vehicle. (**B**) Tumor/liver weight ratio in mice with KPPC tumors treated with vehicle or NXP900 (40mg/kg/dose, once daily). (**C**) Tumor/liver weight ratio in mice with SB1 tumors treated with vehicle or NXP900 (40mg/kg/dose, once daily). (**D**) Schematic depicting the NOD/SCID PDX model established from primary human CCA with subsequent tumor engraftment and treatment with NXP900 or vehicle. (**E**) Dynamic tumor growth rates, expressed as fold change in tumor volume per day, for NXP900 and vehicle-treated mice. Whiskers indicate minimum and maximum values. All data points are shown. (**F**, **G**) Waterfall plots showing the delta tumor volume from the start of treatment to either the first day of statistically significant response or the last day of treatment, whichever occurred first. Only PDX models demonstrating a response to treatment are included. (**G**) IDH-mutated models: IDH1-mutant models PDX 115 and PDX 194 and IDH2-mutant model PDX 276. Statistical analysis was performed with 2-tailed Student t test. Statistical significance: *P < 0.05, **P < 0.01, ***P < 0.001, **** P < 0.0001.

Next, we investigated treatment response to NXP900 in human tumors. Eleven molecularly heterogeneous CCA PDX models (**Suppl. Table 1**) were expanded and implanted into NOD/SCID mice **(Fig. 3D**). We treated tumor bearing mice with vehicle or NXP900 at 40 mg/kg by oral gavage once daily for up to four weeks. Treatment with NXP900 significantly decreased xenograft tumor growth in six out of eleven PDX models (**Fig. 3E**, **Fig. 3F**, **Fig. 3G, Suppl. Fig. 3A**).

Previous studies have observed that IDH-mutant CCA models were especially sensitive to SFK inhibition utilizing the pan-SFK inhibitor dasatinib ^29,30^. Two cell lines with *IDH1* mutations, RBE and SNU-1079, showed similar sensitivity to NXP900 as the other *IDH1/2* wildtype human cell lines (**Fig. 1A**) ^44^. To further explore the sensitivity of the IDH subset specifically, we tested two *IDH1*-mutant models (PDX 115 and PDX 194) and one *IDH2*-mutant model (PDX 276). All three models were sensitive to NXP900 treatment, with more dramatic tumor responses in the *IDH1*-mutant models compared to the sensitive IDH-wildtype models, including reduced tumor burden and significant changes in tumor growth rate compared to vehicle after only 7-9 days of treatment (**Fig. 4E**, **Fig. 4G**).

**Figure 4.**
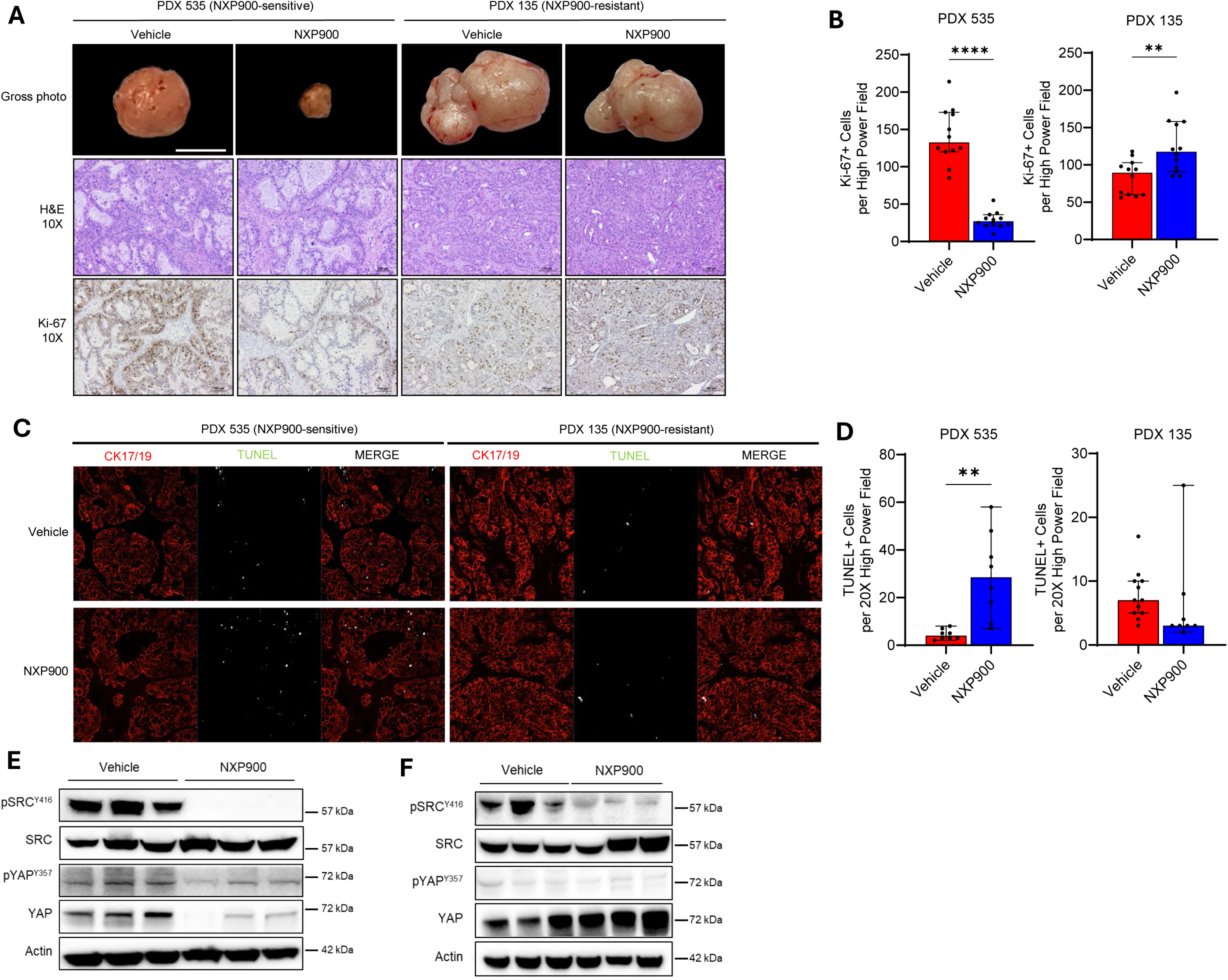
NXP900 downregulates SFK-YAP signaling in vivo. (**A**) Representative liver gross photos and H&E-stained or Ki67 stained tumor sections of PDX 535 (NXP900-sensitive) and PDX 135 (NXP900-resistant) in vehicle- and NXP900-treated mice. Magnification: 10X. Scale bar: 1 cm = 100 μm. (**B**) Quantification of Ki67-positive cells in 4 random high-power fields (HPF) (20X) per section. (**C**) Representative images of TUNEL staining and CK17/19 counterstaining in tumors from vehicle- and NXP900-treated animals. Magnification: 20X (according to methods). Scale bars: 100 μm (according to figures). (**D**) Quantification of TUNEL-positive nuclei of CK17/19 positive cells in at least 4 random HPF (20X) per section. (**E, F**) Tumor lysates from PDX 535 (E) or PDX 135 (F) specimen after treatment with vehicle or NXP900 subjected to immunoblot for pSRCY416, total SRC, pYAPY357, and total YAP. Actin was used as a loading control. Statistical analysis was performed with 2-tailed Student t test. Statistical significance: **P < 0.01, **** P < 0.0001.

Tissue level effects included decreased Ki-67 and increased TUNEL staining, indicating reduced proliferation and increased cell death (**Fig. 4A-D)**. The resistant PDX model 135 did not show a decrease in Ki-67 or increase in TUNEL-positivity. Inhibition of SRC-YAP signaling in tumors of NXP900-treated mice was confirmed in the sensitive model PDX 535 by evaluating the total YAP, SRC and phosphorylated YAP Y357 and SRC Y416 levels by immunoblot (**Fig. 4E**). Although the resistant model PDX 135 did show abrogated SRC activity, no changes in YAP signaling were observed, suggesting a compensating mechanism of YAP Y357 phosphorylation (**Fig. 4F**). NXP900 treatment was well tolerated by the animals and did not result in weight loss compared to vehicle (**Suppl. Fig. 4A**). Serum biochemical analysis noted mild alterations in liver chemistries which mostly remained within normal limits (**Suppl. Fig. 4B**). These data suggest that NXP900 has therapeutic activity against a subset of CCA models *in vivo* and is well tolerated at effective dose levels.

The combination of gemcitabine plus cisplatin is the systemic therapy backbone of first-line therapy for advanced or metastatic biliary tract cancers. Since SRC and YAP activation have been associated with therapeutic resistance, we next assessed whether NXP900 could modulate the sensitivity of CCA cells to the combination of gemcitabine and cisplatin. First, we assessed the *in vitro* effects of combination therapy in HuCCT1 cells and observed that NXP900 plus gemcitabine/cisplatin induced more cell death and a decrease in tumor cell proliferation as compared to NXP900 or gemcitabine/cisplatin alone (**Fig. 5A**, **Fig. 5B**). We then exposed the HuCCT1 and RBE cell lines to increasing concentrations (1 nM - 1 µM) of NXP900 and equimolar concentrations of gemcitabine/cisplatin. The combinatorial effects were evaluated using CalcuSyn software. All combination indices for concentrations between 1 nM and 1 µM were less than 1, indicative of synergy. Strikingly, even in the IDH-mutant RBE cell line, which is expected to be hypersensitive to SFK inhibition as a single agent, we observed synergy with combinatorial therapy. Combination indices in HuCCT1 cells were 0.066, 0.183, 0.427, and 0.501. Combination indices in RBE cells were 0.072, 0.149, 0.466, and 0.885 (**Fig. 5C**).

**Figure 5.**
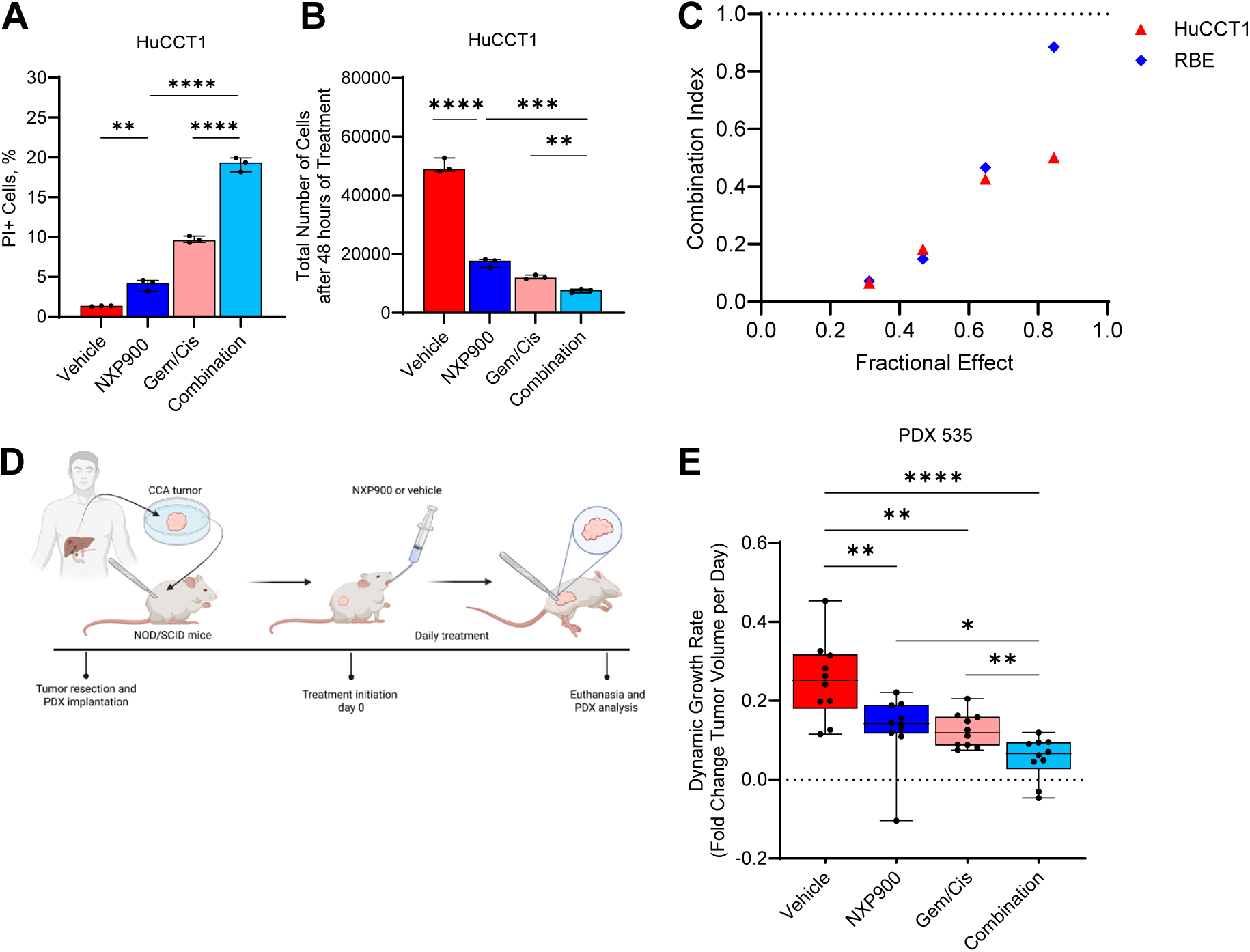
Combination therapy of NXP900 and gemcitabine/cisplatin demonstrates a synergistic interaction. (**A**) Cell death evaluated after staining with propidium iodide in HuCCT1 cells treated with vehicle, NXP900 (20 nM), gemcitabine/cisplatin (100 nM) or combination therapy for 24 hours. (**B**) Cell proliferation evaluated by cell counting after staining with Hoechst 33342 in HuCCT1 cell lines treated with vehicle, NXP900 (20 nM), gemcitabine/cisplatin (100 nM) or combination therapy for 48 hours (*P < 0.05). (**C)** Combination indices (CI) for NXP900 and gemcitabine-cisplatin treatment for 48 hours, as computed by CalcuSyn for HuCCT1 and RBE cells. CI <1 is indicative of synergy. (**D**) Schematic depicting NOD/SCID PDX 535 model tumor development and treatment with vehicle, NXP900 (40 mg/kg), gemcitabine (4 mg/kg)/cisplatin (1 mg/kg) or combination therapy. (**E**) Dynamic tumor growth rates per day (*P < 0.05, **P < 0.01, ****P < 0.0001).

To assess the potential synergistic effect of NXP900 in combination with gemcitabine/cisplatin *in vivo*, we evaluated this therapeutic regimen in the PDX 535 model, which was responsive to both NXP900 monotherapy and gemcitabine/cisplatin (**Fig. 5D)**. The combination of NXP900 with gemcitabine/cisplatin resulted in a significant reduction in tumor growth compared to either monotherapy alone (**Fig. 5E)**. Combination therapy was relatively well tolerated, with weight loss and liver chemistry alterations consistent with the gemcitabine/cisplatin treatment alone (**Suppl. Fig. 4C and 4D**). These data suggest that NXP900 treatment sensitizes CCA to gemcitabine-cisplatin treatment and that combination therapy of NXP900+gemcitabine/cisplatin synergistically induces cell death and inhibits tumor growth.

### Hierarchical all-against-all (HAllA) testing identifies multiomics features associated with primary response to NXP900

To discover molecular biomarkers associated with response, we used a comprehensive multiomic approach, integrating transcriptomics, global protein, and phosphoproteomics analyses of our previously characterized PDX models to identify pathways driving sensitivity and resistance to NXP900 ^45^. We utilized a hierarchical-all-against-all clustering algorithm (HAllA) to assess associations between the abundance value the multiomic features and the drug response values from a total of 8 profiled models treated, including 4 responding and 4 non-responding models (**Fig. 6A, Suppl. Fig. 3B, Suppl. Fig. 3C**) ^46^. Of the 398 significant features, 253 were transcriptomic features, 45 were global protein features, and 100 were phosphoproteomic features.

**Figure 6.**
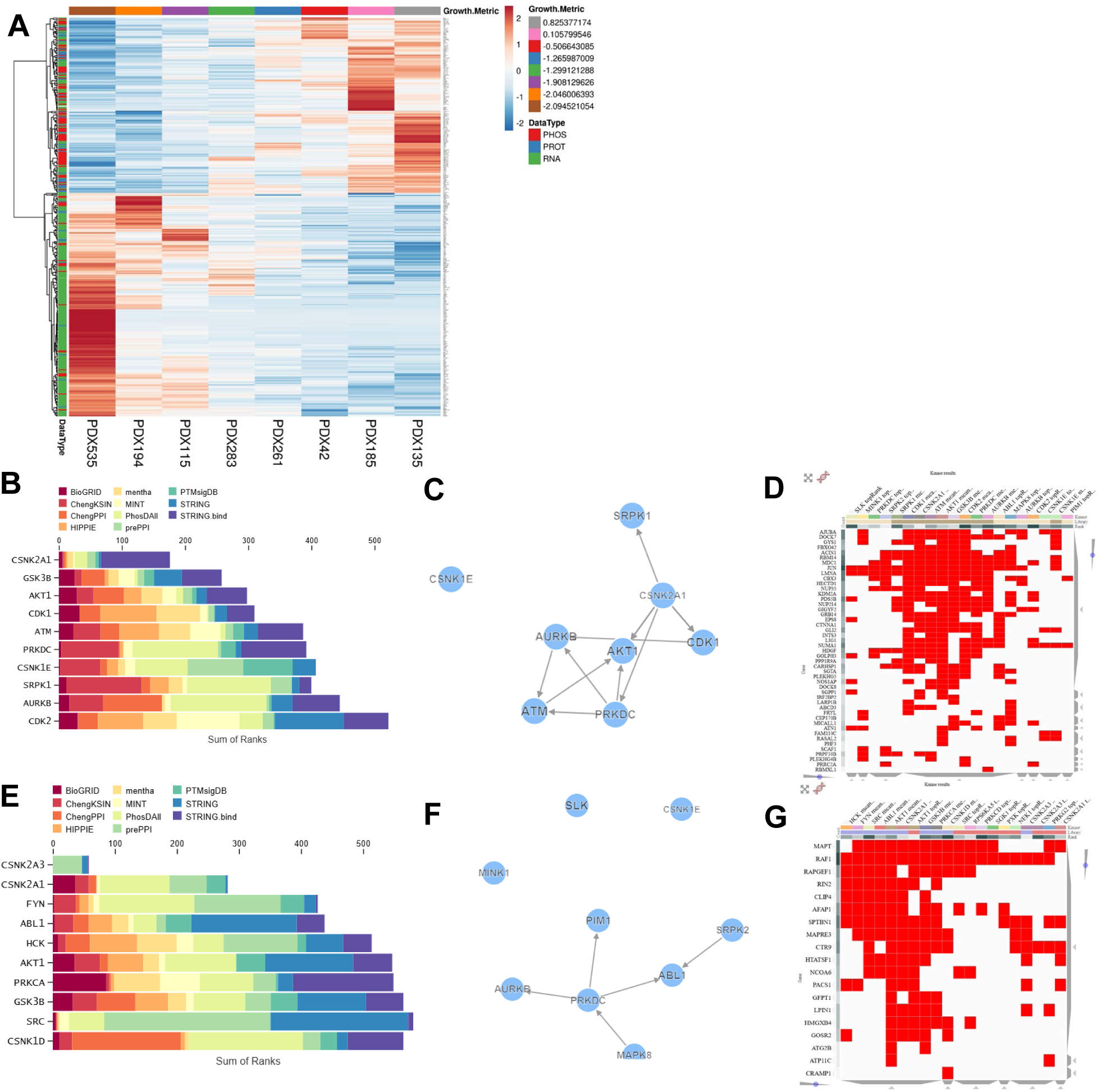
Multiomic determinants of in vivo response. (**A**) Hierarchical-all-against-all clustering algorithm (HAllA)gram of significantly correlated (FDR <0.05) multiomic features associated with the quantitative growth metric in response to NXP900 treatment. Block associations are presented in top-down manner with a descending order of significance. Spearman correlation was used as a similarity metric. Feature values are Zscore normalized across the cohort (increased red, decreased blue). Samples are annotated with the quantitative growth metric. (**B, E**) Kinase enrichment analysis (KEA3) predicting the top kinases enriched in non-responders (B) or responders (E) to NXP900 from 11 different tools, calculated from re-ranking a composite list of kinases by each kinase’s mean integer rank across all libraries containing that kinase. (**C, F**) Kinase network of top kinases enriched in non-responding (C) and responding (F) models from KEA3. (**D, G**) Binary heatmap of substrate evidence for top kinases enriched in non-responding (D) and responding (G) PDX models from KEA3.

Kinase Enrichment Analysis (KEA) using the mean rank score from 11 tools identified several kinases associated with response to NXP900 (**Fig. 6B**, **Fig. 6E**). Models that did not respond to NXP900 demonstrated enrichment of the cell cycle regulators cyclin dependent kinase (CDK) 1 and 2 (**Fig. 6B-D**). As shown by the interacting map (**Fig. 6C**), primary resistance was associated with AKT activity. Notably, kinases that were significantly enriched in the models that did respond included SFKs, including SRC, HCK, and FYN (**Fig. 6E-G**).

To understand the possible clinical utility of NXP900 treatment in CCA, we next examined what proportion of external datasets would respond by mapping external omics data onto our multiomic model. First, the multiomic features were subset to our full cohort of 28 multiomic-profiled PDX models ^45^. Using this approach, we estimated 16 out of 28 (57%) patient tumors may respond to NXP900 (**Fig. 7A**). Second, we assessed an additional internal cohort of 43 multiomic-profiled patient samples, of which 31 were predicted to be responders (72%) (**Fig. 7B**) ^45^. Finally, we analyzed proteogenomic data from the Fudan University (FU)-iCCA cohort ^47^. This large cohort consists of prospectively collected, treatment-naive iCCA samples from China and is reported to be consistent with previous studies. In total, 121 out of 208 samples (58%) were predicted to be sensitive to NXP900 based on their multiomic features (**Fig. 7C**). To determine the possible clinical impact of NXP900 in other tumor types, we mapped multiomic data from 33 distinct cancer types from The Cancer Genome Atlas (TCGA) onto our multiomic model. Up to 50% per were predicted to respond to NXP900, indicating its potential for broad therapeutic applicability (**Fig. 7D**).

**Figure 7.**
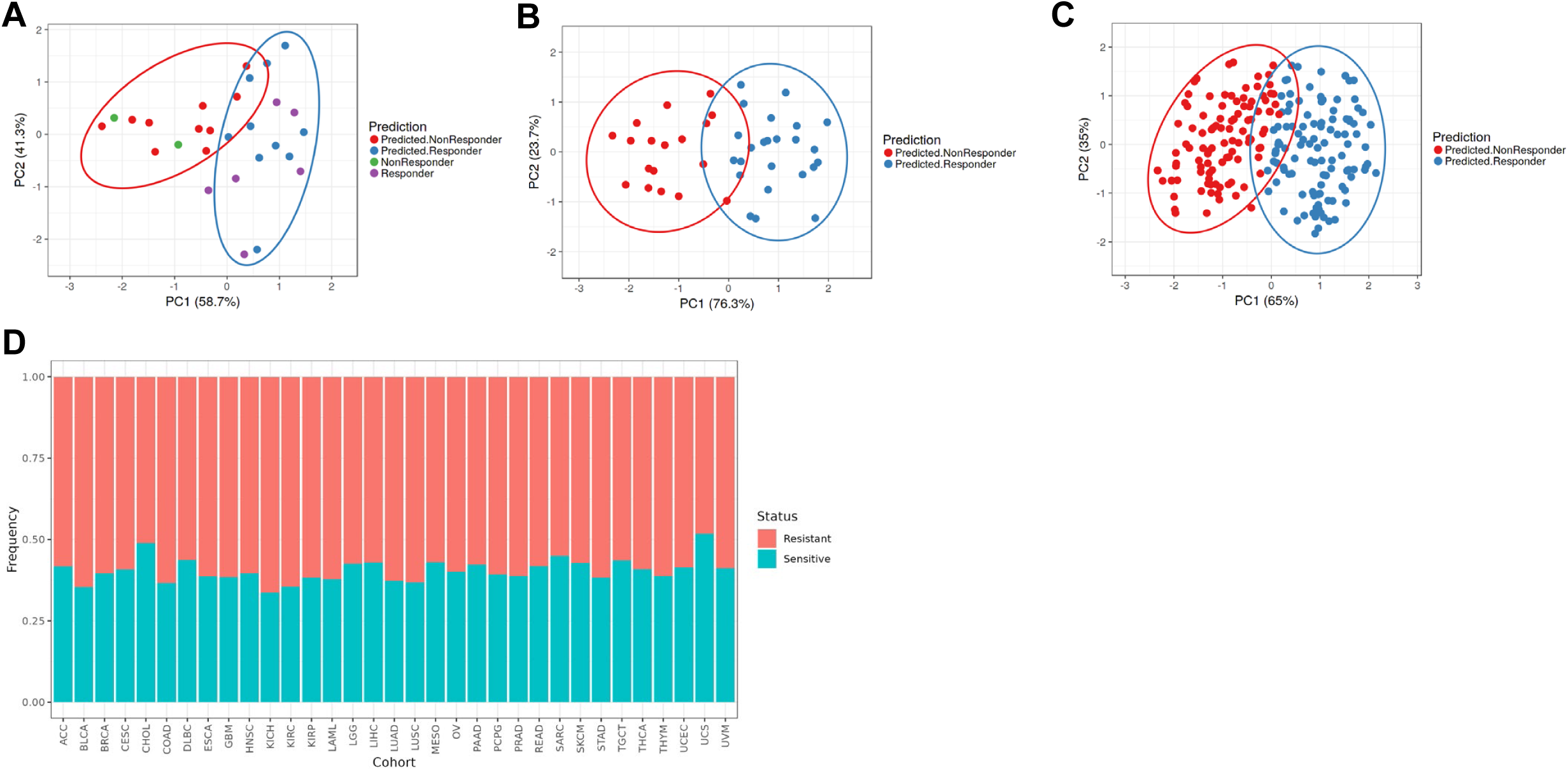
Multiomic profiling predicts clinical utility of NXP900 in CCA and other cancer types. (**A-C**) Principal component (PC)1 vs. PC2 of the principal component analysis (PCA) of an internal cohort of PDX models (A) (responders: 16/28=57%), patients of an internal Mayo Clinic CCA cohort (B) (responders: 31/43=72%), and patients of the external Fudan University-iCCA cohort (C) (responders: 121/208=58%) using significant features associated with response to NXP900. Samples are colored by prediction of response to NXP900 or confirmed response to NXP900 (PDX models only). (**D**) Predicted frequency of tumors per cancer type to respond to NXP900 by mapping 33 distinct cancer types from The Cancer Genome Atlas (TCGA) onto the multiomic response model.

### Acquired mechanisms of resistance to NXP900 are characterized by activated IL13RA/AKT signaling

In order to identify mechanisms of adaptive resistance, we generated cell lines resistant to NXP900 through prolonged exposure to increasing sublethal doses for the CCA cell lines HuCCT1, CCLP1 and SB1 **(Fig. 8A**). Cell viability assays conducted with varying concentrations of NXP900 revealed a substantial increase in the IC50 of resistant cell lines compared to parental lines, with values of 7.9 µM vs. 45 nM in HuCCT1 cells, and 115 µM vs. 15 µM in CCLP1 cells, and 2.6 µM vs. 29.7 µM in SB1 cells, respectively (**Suppl. Fig. 5A**). Notably, the *IDH*-mutant RBE cell line did not tolerate increasing concentrations of drug, and a fully resistant cell line was not able to be developed utilizing this technique. This supports the previous reports suggesting that *IDH*-mutant models have hypersensitivity to SFK inhibition. Resistance was non-reversible, as withdrawal of NXP900 treatment during 10 passages did not decrease the IC50 of resistant HuCCT1 (HuCCT1-R) cells (IC50 4.8 µM vs. 3.6 µM, **Suppl. Fig. 5B**).

**Figure 8.**
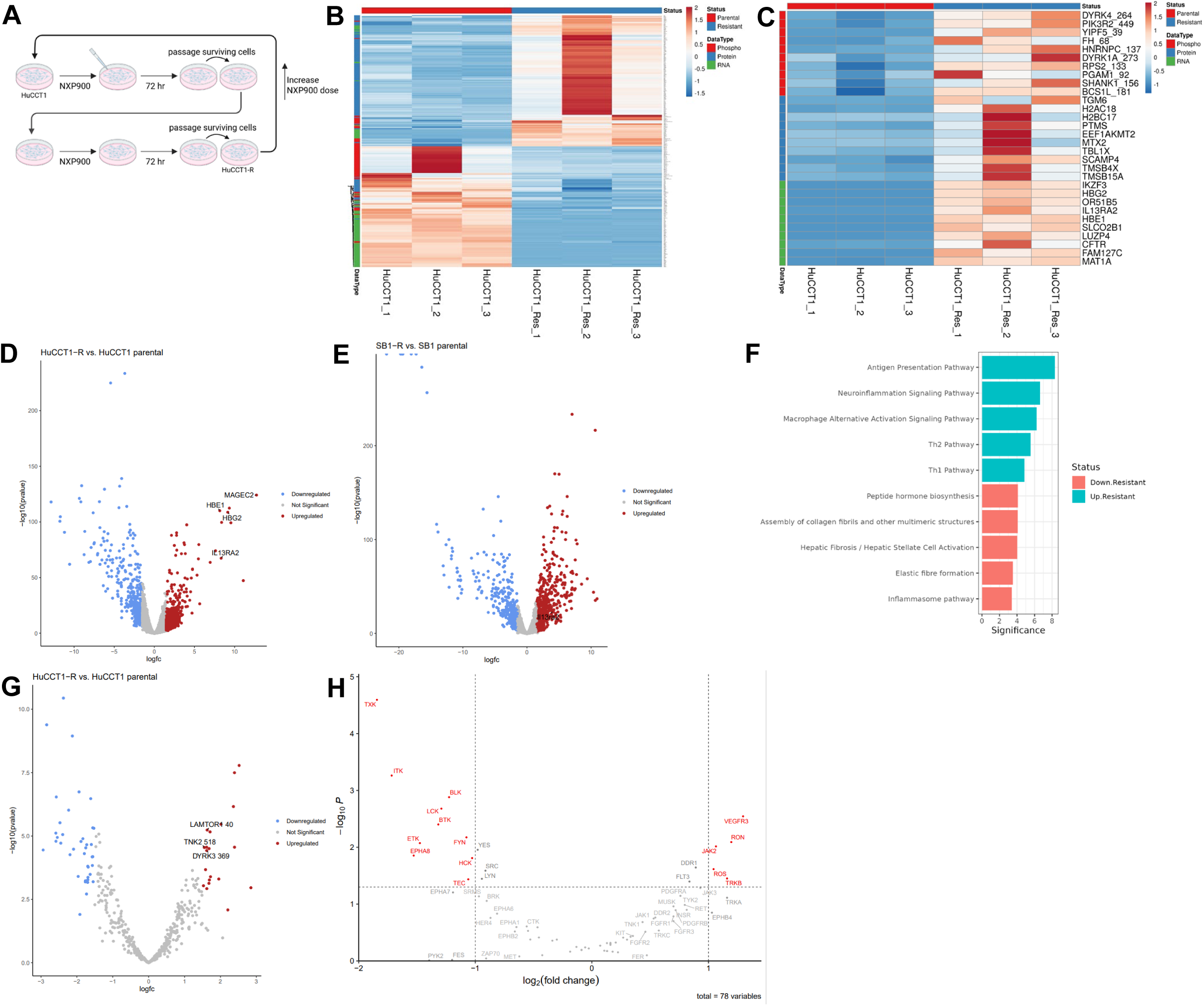
Acquired mechanisms of resistance to NXP900 are characterized by activated IL13RA2 signaling. (**A**) Development of cell lines resistant to NXP900, for example HuCCT1-R. (**B**) Broad characterization of HuCCT1-R cells using RNA-Seq, global proteomics, and phosphoproteomics. (**C**) Top altered differentially expressed genes, proteins and phosphoproteins in HuCCT1-R cells. (**D, E**) Volcano plot comparing RNA-Seq gene expression in parental and resistant HuCCT1 (D) and SB1 cells (E) (3 technical replicates/group). Significantly differentially expressed genes (–1.5 < logFC > 1.5, FDR < 0.05) highlighted blue (downregulated) and red (upregulated) with top genes of interest identified on plot. (**F**) Canonical pathway analysis of the up-and downregulated pathways in HuCCT1-R cells based on RNA-Seq results. Significance is the −log10(PValue) for the pathway. (**G**) Volcano plot comparing phosphoproteomic (pTyr) abundance in HuCCT1 parental and HuCCT1-R cells (3 technical replicates/group). Significantly differentially expressed phosphosites (–1.5 < logFC > 1.5, FDR < 0.05) highlighted blue (downregulated) and red (upregulated) with top phosphoproteins of interest identified on plot. (**H**) Kinase enrichment analysis of Tyr phosphorylation data analysis using a kinase library (https://kinase-library.phosphosite.org/kinase-library/).

We utilized the resistant cell lines and compared these to the parental cell lines to determine multiomic features associated with acquired resistance. Clustering analysis of multiomics showed transcriptomic and proteomic differences between HuCCT1-R and parental sample groups and similarities between the technical replicates (**Fig. 8B**). To investigate acquired resistance mechanisms independent of the cell line used, we performed RNA-Seq analysis on the HuCCT1-R and the SB1-R cell lines. Amongst both cell lines, *IL13RA2*, known to promote tumor cell proliferation, was the top upregulated gene (**Fig. 8C-E)** ^48^. Pathway analysis of the HuCCT1-R cells indicated upregulation of Type 1 and Type 2 T helper responses, neuroinflammation and macrophage signaling pathways, and antigen presentation pathways (**Fig. 8F**). Among the downregulated pathways were hepatic fibrosis activation, elastic fiber formation, and collagen fibrils assembly (**Fig. 8F**).

Phosphoproteomic analysis of the HuCCT1-R cells revealed several tyrosine-phosphorylated proteins that had increased abundance in HuCCT1-R cells compared to HuCCT1 parental cells (**Fig. 8C**, **Fig. 8G)**. To control for changes in total protein abundance, we also performed global proteomics. Amongst the most significantly upregulated phosphosites in HuCCT1-R was TNK2 Y518 (**Fig. 8C**, **Fig. 8G**). TNK2 is known to increase PIP3 levels leading to AKT activation through phosphorylation at T308 and S473. DYRK3 Y369 and LAMTOR1 Y40 were also significantly upregulated, both of which can sustain AKT phosphorylation via mTORC (**Fig. 8C**, **Fig. 8G**). Since short-term treatment of parental cells did not increase these phosphosites, this suggested that these phosphorylation events were associated with a resistance pattern rather than an effect inherent to drug exposure. Kinase enrichment analysis identified differences between the resistant and parental lines (**Fig. 8H**).

Finally, to confirm the stability of the adaptive responses in HuCCT1-R cell line, we repeated multiomic profiling in the HuCCT1-R cells after treatment with NXP900. Comparison of vehicle and NXP900-treated HuCCT1-R cells demonstrated that few genes were significantly altered after treatment, mostly related to downregulation of HLA and heat shock protein (HSP) genes, and that abundance of only one phosphosite, YES1 Y194, was decreased after NXP900 treatment (**Suppl. Fig. 6A-C**). This suggests that these cells are stably altered.

Overall, multiomic profiling of cell lines resistant to NXP900 demonstrated multiple adaptive responses with a strong IL13RA2-AKT signature, suggesting that this signaling axis may be a primary driver of adaptive resistance to SFK inhibition with NXP900.

### Targeting IL13 or AKT signaling can help overcome acquired resistance to NXP900

Given the IL13RA2-AKT signal identified in the CCA cell lines with acquired resistance, we evaluated whether targeting these pathways could resensitize cells to NXP900. The HuCCT1-R cells demonstrated increased proliferation and marked increase in tumor formation in cell line xenograft experiments (**Fig. 9A**). Compared to the parental cells, HuCCT1-R cells demonstrated a higher tumor engraftment rate (75% vs. 35%) and greater tumor size (median weight 632.5 mg vs. 66.4 mg) (**Fig. 9B, 9C**). Knockdown of *IL13RA2* in HuCCT1-R cells significantly reduced cell proliferation and resensitized HuCCT1-R cells to NXP900 treatment (**Fig. 9D**, **Suppl. Fig 7)**. Because IL13RA2 can activate AKT signaling, we evaluated AKT phosphorylation and observed that *IL13RA2* knockdown resulted in a parallel downregulation in AKT activation (**Fig. 9E**) ^49,50^. We then explored targeting AKT directly in the resistant cell line utilizing the FDA-approved pan-AKT inhibitor capivasertib. Similar to the results with *IL13RA2* downregulation, capivasertib in combination with NXP900 reduced cellular proliferation and increased cell death consistent with overcoming acquired resistance (**Fig. 9F**). Taken together, these data support a model where adaptive resistance to the SFK inhibitor NXP900 is accomplished by upregulation of an IL13RA2-AKT signaling axis which can be directly targeted to improve or restore therapeutic response.

**Figure 9.**
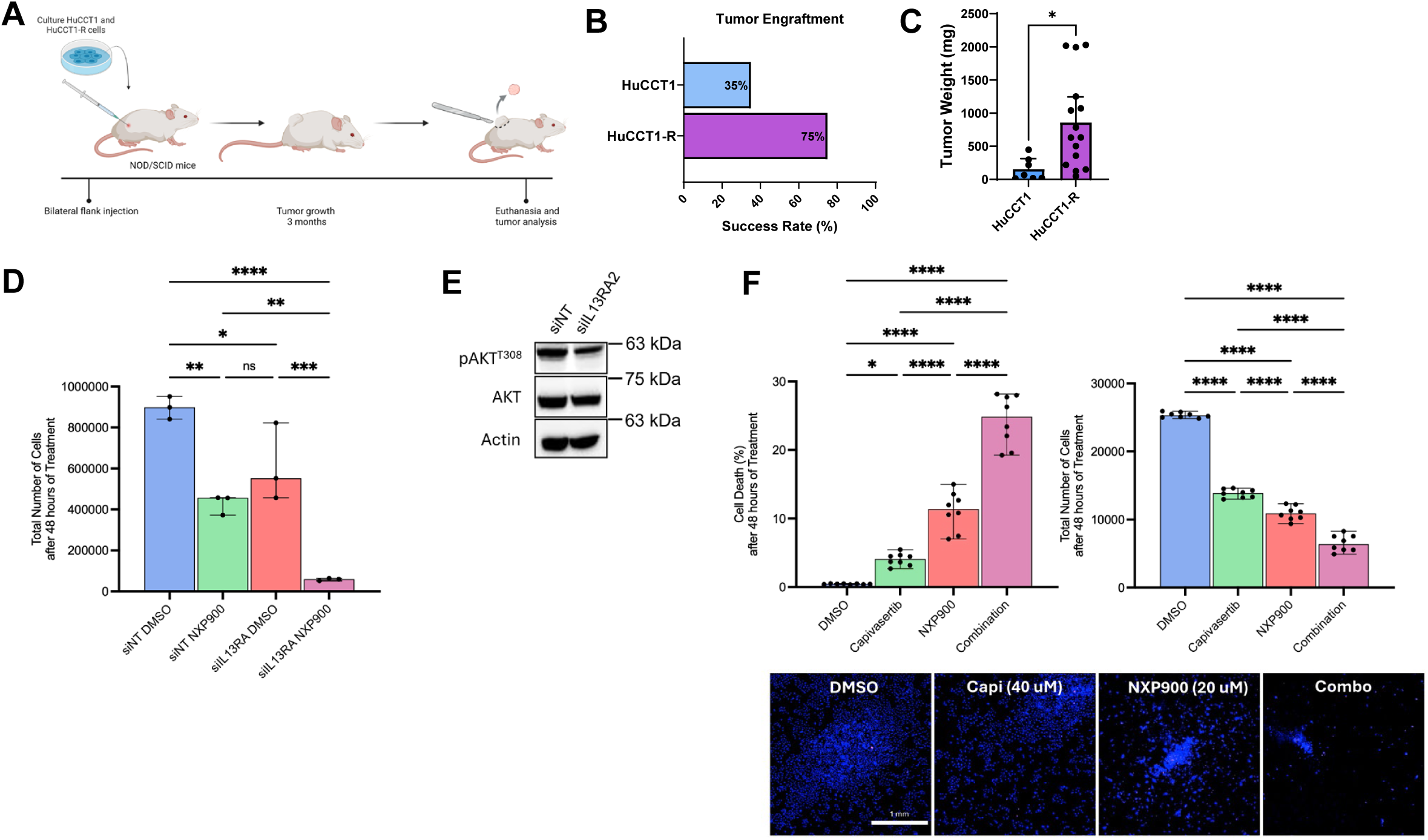
Acquired resistance to NXP900 is overcome by targeting IL13 or AKT signaling. (**A**) Schematic depicting a NOD/SCID mice xenotransplantation model of HuCCT1 and HuCCT1-R cells by bilateral subcutaneous flank injections. (**B**) Tumor engraftment success rates of HuCCT1 and HuCCT1-R injections. (**C**) Tumor weights (mg) three months post-injection. (**D**) Cell proliferation evaluated by cell counting after staining with Hoechst 33342 in HuCCT1-R cell lines transfected with targeting and nontargeting siRNA for IL13RA2, treated with vehicle (DMSO) or NXP900 for 48 hours (*P < 0.05). (**E**) Cell lysates from the HuCCT1-R cell line with siIL13RA2 or siNT for 48 hours subjected to immunoblot for pAKT T308 and total AKT. Actin was used as a loading control. (**F**) Cell death and proliferation in HuCCT1-R cells treated with either NXP900 (20 µM), capivasertib (40 µM), or a combination, for 48 hours. Representative images showing the total number of cells after staining with Hoechst 33342 following the four different treatments.

## DISCUSSION

This study presents a comprehensive analysis of SFK inhibition, utilizing the novel SFK OFF inhibitor NXP900, in multiple preclinical models of CCA. Evaluation identified treatment induced molecular alterations, multiomic biomarkers of primary response, and mechanisms of adaptive resistance. Specifically, the major findings of this study indicate that: (1) NXP900 provides a novel therapeutic strategy to target YAP activity and leads to cell death in multiple preclinical models of CCA; (2) patient-derived tumors with *IDH-*mutations are hypersensitive to SFK inhibition with NXP900; (3) primary resistance to NXP900 is associated with activated AKT; (4) activation of an IL13RA2-AKT pathway is a key component of acquired resistance to NXP900 and can be overcome by IL13 or AKT targeting. These findings are discussed in greater detail below.

SFKs play a role in regulating normal and oncogenic processes, including proliferation and survival ^51^. Despite SFK activity in many tumors, no SFK inhibitors have received approval for treatment of solid tumors yet ^52,53^. SFK inhibitors like dasatinib and bosutinib have shown modest clinical benefit and unfavorable treatment toxicity, possibly due to off-target effects ^53,54^. These inhibitors impair kinase catalytic activity but bind to the kinase in its open conformation, thus inducing scaffolding functions and interaction with other proteins. However, structural analysis of the NXP900 revealed a distinct mechanism in which it locks SRC in its native inactive ‘OFF’ conformation, inhibiting both kinase activity and scaffolding functions. In addition to this unique mode of inhibition differentiating NXP900 from other SFK inhibitors, NXP900 demonstrates a highly selective profile with minimal inhibition of kinases outside the SFK family ^33,55^. Moreover, compared with dasatinib, NXP900 demonstrates superior potency of SRC inhibition at lower concentrations, and shows a more robust antitumor effect as well as delayed tumor relapse in preclinical models ^33,55^. Previous *in vitro* investigations of NXP900 have demonstrated inhibited tumor cell proliferation and induced cell cycle arrest following treatment ^33,55^. These activities translated into *in vivo* efficacy in various murine models including breast cancer, enzalutamide-resistant prostate cancer bone metastasis, ovarian cancer, esophageal cancer, and head and neck carcinomas ^56–58^. Of particular significance is the initiation of the Phase 1 clinical trial of NXP900 (NCT05873686). Phase 1a of this clinical trial assessed the safety, tolerability and pharmacokinetic properties of NXP900 in patients with advanced, metastatic and/or progressive solid tumors and demonstrated an acceptable safety profile along with a robust pharmacodynamic response at doses ≥ 150 mg/day ^59^. A subsequent Phase 1b program was initiated to evaluate the clinical activity of NXP900 as a single agent in patients with advanced solid tumors with YES1 amplifications or Hippo pathway alterations. This marks an exciting opportunity to evaluate the translation of preclinical efficacy into clinical settings.

Altering YAP signaling remains an area of interest because YAP is activated in several cancers yet clinically available direct inhibitors are lacking ^60,61^. Furthermore, targeting YAP through the Hippo pathway has been proven difficult, as only a small subset of patients with CCA have identifiable mutations in the Hippo pathway components, such as a *NF2* mutation (2.6%) or *SAV1* deletion (5.3%) ^9^. Finally, YAP activation is now commonly recognized as a mechanism of acquired resistance in the setting of chemotherapy and targeted therapeutics, including KRAS inhibition ^62,63^. Therefore, our finding that NXP900 inhibits YAP activity represents a significant advancement with potential broad clinical implications for CCA and other malignancies.

PDX models harboring an IDH1 mutation exhibited hypersensitivity to NXP900 treatment, consistent with previous findings ^29,30^. Sensitivity to SFK inhibitors such as the pan-SFK inhibitor dasatinib in IDH1 mutant-tumors, has been linked to a signaling axis involving the inhibition of the tumor-suppressive membrane-associated guanylate kinase, WW, and PDZ domain containing 1 (MAGI1) - protein phosphatase 2A (PP2A) complex by SRC to activate p70 S6 kinase (S6K) and ribosomal protein S6 (S6) signaling ^29,30^. Current *IDH1*-targeting therapies such as ivosidenib have demonstrated limited efficacy, primarily inducing stable disease rather than significant tumor regression, and are often compromised by acquired resistance ^31,32^. In contrast, the rapid tumor shrinkage observed in our *IDH1*-mutated PDX models following only a few days of NXP900 treatment suggests that SFK inhibition may be a promising therapeutic strategy for patients with *IDH1*-mutated CCA beyond standard *IDH1*-targeting therapies. The sensitivity of an *IDH2*-mutated model is of particular clinical relevance, given the current lack of approved therapies specifically targeting *IDH2* mutations in CCA ^64^.

Comprehensive multiomic profiling of eight CCA PDX models revealed that activation of the AKT pathway is associated with primary resistance to NXP900. This was further supported by resistance observed in the syngeneic SB1 model, which is driven by constitutively active myr-AKT. In contrast, sensitivity to NXP900 correlated with SFK activity, consistent with the drug’s proposed mechanism. To assess the broader clinical relevance, we applied our integrated multiomic response model to additional cohorts, predicting NXP900 sensitivity in 57–72% of CCA tumors and up to 50% across other cancer types. These findings suggest that a significant subset of patients with CCA and other malignancies may benefit from NXP900 treatment. The utility of this response model will need to be tested in a prospective clinical trial but suggests that study design will be important to optimize the patient population most likely to benefit from this drug.

Acquired resistance to NXP900 was characterized by upregulation of IL13RA2-AKT signaling. *IL13RA2* is implicated in promoting cancer cell proliferation, survival and metastatic potential ^49,65^. By silencing *IL13RA2*, we were able to effectively reduce tumor cell proliferation *in vitro*. Previous studies have demonstrated that IL13 signaling in cancer cells activates MTOR, FAK, SFK, AKT, and ERK1/2 pathways ^49,50^. Dual inhibition with NXP900 and the pan-AKT inhibitor, capivasertib, resensitized resistant cells to NXP900. Inhibiting either IL13 or AKT signaling presents a potential clinical strategy to overcome resistance to NXP900. In normal tissues, the expression of *IL13RA2* is restricted to the testis, making it ideal targets for targeted therapies such as CAR-T cell and T-cell receptor therapy ^66,67^. A recent phase 1 trial on IL-13Rα2-targeting CAR-T cells with locoregional therapy showed safety and clinical activity in a subset of patients ^68^. Capivasertib is already approved by the FDA in combination with fulvestrant for HR-positive, HER2-negative locally advanced or metastatic breast cancer with AKT pathway-alterations (PIK3CA, AKT1, or PTEN) ^69^. Future studies should investigate the efficacy of combined NXP900 and capivasertib treatment to either prevent resistance or to resensitize resistant tumors as a strategy to improve treatment outcomes.

Taken together, our study evaluated treatment induced molecular alterations of SFK inhibition with NXP900 and identified biomarkers of response and mechanisms to overcome resistance to SFK inhibition. Finally, the novel SFK ‘OFF’ inhibitor NXP900 could be a new therapeutic option to inhibit YAP in patients with CCA and other malignancies.

## METHODS

### Cell culture

The human CCA cell lines HuCCT1 (FBXW7ΔF/KRAS/TP53), RBE (IDH1/KRAS), (KRAS/TP53), SNU-1079 (IDH1/FGFR2/CCND2) and CCLP1 (CTNNB1/TP53/SMARCA4), and murine CCA cell lines SB1 (myr-AKT/YAP^S127A^) and KPPC (KRAS^G12D^/TP53-/-) were cultured in DMEM supplemented with 10% FBS, 1% penicillin-streptomycin and 0.2% primocin and maintained in a humidified atmosphere at 37 °C in the presence of 5% CO_2_ ^42^. Because of cell density regulation of the Hippo pathway, cell culture experiments were performed at near confluence (∼80%). Clones of CCA cell lines resistant to NXP900 were generated through a process of slowly escalating exposure to NXP900 for over 6 months. Resistant cells were maintained in culture in the presence of the last well-tolerated concentration of NXP900 (i.e., 100 nM for HuCCT1, 1 μM for CCLP1 and SB1).

### Compounds and antibodies

NXP900 (Nuvectis Pharma), capivasertib (MedChemExpress), gemcitabine (Novaplus) and cisplatin (West-Ward Pharmaceuticals Corp) were added to cells at final concentrations from 0.001 nM – 1 mM. The following primary antibodies were used for immunoblot analysis: actin (sc-8432) (Santa Cruz Biotechnology), YAP (sc-101199) (Santa Cruz Biotechnology), phospho YAP (Y357) (ab62751) (abcam), phospho YAP (S127) (4911) (Cell Signaling Technology), Src (2110) (Cell Signaling Technology), phospho Src Family (Y416) (2101) (Cell Signaling Technology), LATS1 (3477) (Cell Signaling Technology), phospho LATS1 (S909) (9157) (Cell Signaling Technology), phospho LATS1 (T1079) (8654) (Cell Signaling Technology), Akt (pan) (4691) (Cell Signaling Technology), phospho Akt (T308) (9275) (Cell Signaling Technology), and IL-13RA2/CD213a2 (85677) (Cell Signaling Technology). The following primary antibodies were used for IHC: YAP (sc-101199) (Santa Cruz Biotechnology) and Ki-67 (12202) (Cell Signaling Technology).

### Cell viability, cell death, and apoptosis assays

For in vitro studies, NXP900 compounds were weighed on a calibrated balance and dissolved in 100% DMSO. To determine the half maximal inhibitory concentration (IC50) for each cell line, cells were seeded in triplicate in 96-well plates at a density of 5.0 x 10^3^ cells per well. After incubation for 24 hours, compound dilutions in 5- or 10-fold steps in complete media were added to the wells. After 48 hours of treatment, CellTiter-Glo® (CTG) (Promega) solution to was added to the wells and luminescence was determined using a BioTek Synergy^TM^ H1 microplate reader. IC50 curves were generated/plotted using GraphPad Prism version 9.1.0 (San Diego, CA, USA). For combination studies, interactions of combinatorial effects were measured by cell viability and expressed as the Combination Index (CI) with CalcuSyn software version 2.11 (Biosoft, Cambridge, UK). Fractional effect (fa) was calculated as: fa=(1-g [Growth inhibition in %])/100.

Cell death was determined after exposure to compounds for 48 hours. Cells were then incubated with Propidium Iodide (PI) 2 µg/mL and Hoechst 33342 (Life Technologies Corporation, Eugene, OR, USA) 5 µg/mL for 30 minutes at 37°C and imaged using a Celigo Image Cytometer (Nexcelom Bioscience, Lawrence, MA, USA). Cell death was plotted using GraphPad Prism version 9.5.1 (San Diego, CA, USA).

Apoptotic effects were measured after 24 hours of treatment using Caspase-Glo 3/7 assay reagent from Promega (Madison, WI, USA). Plates were covered and placed on a shaker for 45 minutes to ensure mixture of cells with reagent. Fluorescence was measured using BioTek Synergy^TM^ H1 microplate reader.

### Quantitative reverse transcription PCR (RT-qPCR)

Tumor samples were disrupted and homogenized in Trizol (Invitrogen) using a rotor-stator homogenizer. PCR-mRNA was isolated from cells and PDX tumor samples using TRIzol Reagent followed by isopropanol precipitation, per manufacturer protocol. RNA concentration was measured using a spectrophotometer (NanoDrop One, Thermo Scientific). Reverse transcription was performed using Moloney murine leukemia virus reverse transcriptase and random primers (Life Technologies). After reverse transcription, cDNA was analyzed by real-time PCR (LightCycler 480 II, Roche Diagnostics) to quantify the target genes; SYBR Green (Roche Diagnostics) was used as the fluorophore. Expression was normalized to 18 S and relative quantification performed according to the 2−ΔCT or 2−ΔΔCT method as previously described ^70^. Data are reported as fold expression compared to calibrator as geometric mean and geometric standard deviation of expression relative to calibrator. Technical replicates were completed for each run and a minimum of three biologic replicates completed for each condition/cell line. The primers used are listed in **Suppl. Table 2**.

### Immunoblot analysis

Whole-cell lysates were collected by adding ice-cold lysis buffer containing protease inhibitors, phosphatase inhibitors and 1% phenylmethylsulfonyl fluoride (PMSF). Cells were scraped to collect cells, vortexed and lysed on ice for approximately 20 minutes. Cells were then subjected to centrifugation at 12,000 ×*g* for 15 minutes at 4°C. Supernatants were transferred to a clean tube to remove cellular debris. Protein concentrations were quantified by Bradford assays (Sigma-Aldrich). For immunoblot analysis, proteins were resolved by SDS-PAGE and transferred to nitrocellulose membranes. Membranes were incubated with primary antibodies (1:1000 dilution) at 4°C overnight in 5% BSA-TBS Tween. After incubation, membranes were washed for 30 minutes in TBS Tween. Horseradish peroxidase–conjugated secondary antibodies against mouse, rabbit, or goat as indicated (Santa Cruz Biotechnology) were then added to membrane at a concentration of 1:5000 and incubated for 1 hour at room temperature. Immunoblots were visualized with enhanced chemiluminescence (ECL) or ECL Prime (GE Healthcare Life Sciences).

### Immunohistochemistry

Tumor tissues were fixed in 10% formaldehyde, paraffin-embedded, and sectioned onto slides by standard protocols. Sections were deparaffinized, hydrated and incubated with primary antibodies listed above for Ki-67 (1:150) at 4°C overnight. Slides were then incubated with secondary antibodies diluted in blocking buffer for 1 hour at room temperature in the dark. Positive cells were counted manually using ImageJ in at least 4 random fields (20X) per tumor and 3 tumors per treatment group.

### TUNEL staining

The fluorescent TUNEL assay (*in situ* cell death detection kit, Roche) was utilized on tissue sections. Sections were deparaffinized and hydrated. The TUNEL assay was then performed using the manufacturer’s protocol and incubated with CK17/19 primary antibody (1:100). The Vector® TrueVIEW® Autofluorescence Quenching Kit was used to quench autofluorescence. Slides were incubated with DAPI staining solution (1:25,000 in 1X PBS) for approximately 10 minutes and mounted with VECTASHIELD Vibrance® Antifade Mounting Medium. Slides were analyzed by fluorescent confocal microscopy (LSM 780, Zeiss). Dead cells were quantified by counting TUNEL-positive nuclei of CK17/19 positive cells in at least 4 random high-power fields per section (20X).

### RNA interference and transfection

IL13RA2 was transiently knocked down in the HuCCT1-resistant cell line with validated siRNAs (Dharmacon). Cells were grown in 6-well plates and transfected using Lipofectamine RNAiMAX Transfection Reagent (Invitrogen) following the manufacturer’s instructions. After 24 hours of incubation, cells were treated with either vehicle or NXP900 (2 µM) for 48 hours. The efficiency and specificity of knockdown were assayed by immunoblot.

### *In vivo* patient-derived xenograft generation and treatment

Our group has previously successfully developed several patient-derived xenograft (PDX) models from surgically resected tissue and analyzed these for genomic and proteomic differences in addition to their clinical and translational relevance ^45,71–73^. In the current study, eleven different PDX models (**Suppl. Table 1**) were xenotransplanted into the flanks of NOD/SCID mice, aged 6-8 weeks. Our study examined male and female animals, as similar findings are reported for both sexes, although female mice were preferred for PDX studies due to practical housing considerations and the absence of significant gender-based differences in tumor engraftment or treatment response ^74^. Treatment studies were initiated when tumors reached palpable size (>125 mm^3^). Tumor bearing mice were randomized 1:1 into treatment and control arms, with 5 to 10 mice per arm. Tumor volume (length x width^2^) and animal weight were measured at baseline and subsequently twice weekly using calipers. Change in tumor volume was determined as difference between baseline measurements and endpoint measurements. NXP900 compounds were prepared at a concentration of 10 µg/µL in vehicle (3 mmol/L sodium citrate buffer pH 3.0). Mice were treated with 100µL of NXP900 (40 mg/kg) once daily by oral gavage, gemcitabine (4 mg/kg)-cisplatin (1 mg/kg) twice weekly by intraperitoneal injection, combination, or vehicle for up to four weeks. At the end of the treatment period, animals were sacrificed, and tumors were extracted for analyses. Blood was collected for measurement of serum alkaline phosphatase (ALP), alanine aminotransferase (ALT), albumin (ALB), and blood urea nitrogen (BUN) (Vetscan V2, Abaxis).

### *In vivo* syngeneic orthotopic murine models of CCA

Murine SB1 or KPPC CCA cells were harvested and washed in DMEM. Male C57BL/6J mice from Jackson Labs were anesthetized using 1.5–3% isoflurane. Male mice were used for implantation to avoid cross-sex immunogenicity because these murine cell lines were derived from tumors grown in male mice. Under deep anesthesia, the abdominal cavity was opened by a 1 cm incision below the xiphoid process. A sterile cotton tipped applicator was used to expose the superomedial aspect of the lateral lobe of the liver. Using a 27-gauge needle, 30 μL of 50% DMEM and 50% Matrigel containing 7.5 × 10^5^ cells was injected orthotopically into the left lateral lobe of the mouse liver. The cotton tipped applicator was held over the injection site to prevent cell leakage and encourage hemostasis. Subsequently, the abdominal wall and skin were closed in separate layers with absorbable 4-0 Vicryl suture material.

Treatments were initiated 7 days after KPPC cell implantation and 14 days after SB1 cell implantation, given differences in the growth rates of these models ^42,43^. Tumor presence was confirmed using the Scintica Prospect T2 small animal ultrasound prior to treatment start. Mice were randomly and equally assigned to vehicle control or treatment groups. KPPC mice were treated for 1 week while SB1 mice were treated for 2 weeks. The control group received 100 µL of vehicle (3 mmol/L sodium citrate buffer pH 3.0) once daily by oral gavage. The treatment group received 100 μl NXP900 at a concentration of 10 µg/µL in vehicle (40 mg/kg) once daily by oral gavage. At the end of the treatment period, animals were sacrificed, and blood was collected for measurement of serum ALP, alanine ALT, ALB, and BUN (Vetscan V2, Abaxis).

### Isolation of tumor-infiltrating immune cells and flow analysis

Upon excision, tumors were dissociated with gentleMACS Octo Dissociator (Miltenyi) using the mouse tumor dissociation kit (Miltenyi 130-096-730) for tumor samples according to the manufacturer’s protocol and as previously described ^75^. CD45 + cells were isolated by CD45 (TIL) mouse microbeads (Miltenyi). Cells were stained and data were acquired on a Miltenyi MACSQuant Analyzer 10 optical bench flow cytometer as previously described ^75^. All antibodies were used following the manufacturer’s recommendation. Fluorescence Minus One controls were used for each independent experiment to establish gates. For intracellular staining of granzyme B, FoxP3, and CD206, cells were stained using the intracellular staining kit (Miltenyi). Apoptosis staining was performed using the APC Annexin V Apoptosis Detection Kit with 7-AAD (BioLegend). Data analysis was conducted using FlowJo (TreeStar). Forward scatter and side scatter were used to exclude cell debris and doublets. Fixable Viability Stain 510 (BD Horizon) was used to exclude dead cells.

### *In vivo* flank implantation model

A NOD/SCID mice xenotransplantation model was developed by bilateral subcutaneous flank injections of HuCCT1 or HuCCT1-R cells (100 μL of 50% DMEM and 50% Matrigel containing 1 × 10^6^ cells) into the left and right flank areas of each mouse. Three months after tumor cell injection, mice were sacrificed, tumors were extracted, and tumor weights were collected.

### RNA isolation for RNA-Seq analysis

Samples of human CCA cell lines were lysed and homogenized in TRIzol Reagent (Invitrogen) per manufacturer protocol with additional on-column DNase treatment (Qiagen #1023460) and RNA wash with Buffer RPE and Buffer RWT (Qiagen).

### Illumina Stranded mRNA library prep with Illlumina NovaSeq 6000 Sequencing

RNA libraries were prepared using 200 ng of total RNA according to the manufacturer’s instructions for the Illumina Stranded mRNA Ligation Sample Prep Kit (Illumina, San Diego, CA). The concentration and size distribution of the completed libraries were determined using an Agilent Bioanalyzer DNA 1000 chip (Santa Clara, CA) and Qubit fluorometry (Invitrogen, Carlsbad, CA). Libraries were sequenced at 50M fragment reads per sample following Illumina’s standard protocol using the Illumina NovaSeq™ 6000 S4 flow cell. S4 flow cells were sequenced as 100 × 2 paired end reads using NovaSeq S4 sequencing kit and NovaSeq Control Software v1.8.0. Base-calling was performed using Illumina’s RTA version 3.4.4.

### Protein extraction and tandem mass tag labeling for proteomics

Proteins were extracted from human CCA cell lines. After trypsinization, cells were subjected to centrifugation at 700 ×*g* for 5 minutes at 4°C. Cells were lysed in lysis buffer (20 mM HEPES pH 8, 8 M urea, 1 mM sodium orthovanadate, 2.5 mM sodium pyrophosphate, 1 mM β-glycerophosphate), and sonicated using a tip sonicator (Branson, SFX 550). Proteins were reduced with dithiothreitol (Sigma, D9163), and subsequently alkylated with iodoacetamide (Sigma, I1149). Protein extracts were diluted with 20 mM HEPEPS to a final concentration of ∼1M urea and TPCK-treated trypsin (Worthington, LS003744) was added with 1:50 enzyme to protein ratio. After overnight incubation at 37 °C, protein digests were acidified and subjected to clean-up using Sep-Pak C_18_ cartridges (Waters). For tandem mass tag labeling, peptides were reconstituted in 100 mM triethyl ammonium bicarbonate. Each sample was labeled using tandem mass tag reagents (Thermo, A52045) following the manufacturer’s instructions. The pooled sample was desalted using Sep-Pak C_18_ cartridges and subjected to phosphopeptide enrichment.

### Immunoaffinity purification of phosphotyrosine peptides

Around 2 mg of lyophilized TMT-labeled peptides were reconstituted in 1.4 mL of IAP buffer (50 mM MOPS pH 7.2, 10 mM sodium phosphate, 50 mM NaCl) followed by incubation with anti-phosphotyrosine antibody beads (PTMScan HS p-Tyr-1000, 38572, Cell Signaling Technology) at 4 °C for 1 hr. After incubation, phosphotyrosine peptides and the antibody complex were washed three times with wash buffer and then twice with water. Elution of phosphopeptides was performed with 0.15% TFA at room temperature. Eluents were dried for LC-MS/MS analysis.

### Mass spectrometry data acquisition and analysis

LC-MS/MS data of TMT labeled peptides were acquired on Orbitrap Exploris 480 mass spectrometer (Thermo Scientific, San Jose, CA) coupled to Ultimate 3000 liquid chromatography system (Thermo Scientific, San Jose, CA). The peptides were loaded onto a trap column (Optima C_18_, 2cm x 100µm, 100 Å) at a flow rate of 8 µl/min and separated on an analytical column (75 µm x 25 cm, C_18_, 1.7 µm, IonOpticks, AUR3-25075C18-TS) at a flow rate of 250 nL/min. Solvent A (0.1% formic acid in water) and solvent B (0.1% formic acid in acetonitrile) were used for generating gradient (3-30%) over 100 min. Mass spectrometry data were acquired in data-dependent acquisition mode with cycle time of 2 sec. MS scan and MS/MS scan were acquired with resolution of 120,000 and 45,000, respectively. Precursor ions were isolated under 1.2 m/z isolation width and fragmented with normalized collision energy (NCE) of 34.

### Multiomic data processing by differential expression analysis

Relative protein intensities were normalized before performing differential expression by making the counts a ratio of the sample’s total (phospho)proteomic counts divided by a scaling factor. Next, edgeR was used to generate a negative binomial model based on the counts and estimates dispersion across genes ^76^. P-values were calculated using a quantile-adjusted conditional maximum likelihood (qCML) method for experiments with single factor.

### Prediction of primary sensitivity and resistance to NXP900 using multiomic data of PDX models

To explain response to NXP900, the relevance of each factor from all multiomic data (transcriptomics, proteomics, and phosphoproteomics) of each previously profiled PDX model was calculated by taking the inverse correlation between the drug response values (relative growth compared to vehicle only treatment) and the factor values for the subset of validated samples. Drug response values at last day of treatment were calculated as Inhibitory Indices Values <0 indicate an inhibitory effect while values >0 indicate a growth effect.

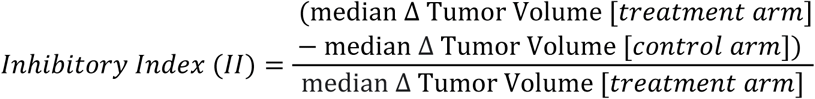

To determine the likelihood of sensitivity for additional PDX samples, a sensitivity score was defined as the summation across all model factors (i) of the absolute value of the product of the NXP900 drug’s correlation with each factor based on the validated samples (Fc) and the factor value of the query sample (Fv) divided by the square root of the number of factors.

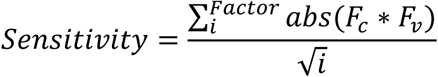

Additionally, the validated PDX models were used to identify features across all multiomics by comparing the drug response values to the quantitative abundance values of each feature using a novel hierarchical framework, hierarchical-all-against-all clustering algorithm (HAllA) ^46^. The significant features were subset into features increased in responding models and features increased in non-responding models. HAllA outputs a correlation coefficient, p-value, and adjusted p-value (FDR) for each x-y feature pair. To identify signaling driving resistance, phosphoproteomic features that were increased in resistant models were input into a kinase enrichment analysis (KEA3), identifying the top ranked kinases from 11 different tools and generating maps of interacting kinases. To identify pathways driving resistance, pathways were identified from the resistance-specific features using GSEA pre-ranked analysis using the HAllA calculated false discovery rate (FDR) and the sign of correlation.

### Statistics

Statistical analyses were performed using two-tailed Student *t* test or Mann–Whitney U test and GraphPad Prism 10 Software (GraphPad Software Inc.). Paired *t* test was used because data were normally distributed. All data are presented as mean ± SD and *P*-values less than 0.05 were considered significant.

## Study approval

Mayo Clinic, Rochester, MN, USA, has an ongoing IRB-approved protocol for the collection and xenotransplantation of resected biliary tract tumors (19-012104 and 70703). All in vivo studies were approved by the Institutional Animal Care and Use Committee at Mayo Clinic, Rochester, MN, USA (A00003954-18-R21 and A00004672-19-R22).

## Data availability

Values for all data points in graphs are reported in the Supporting Data Values file.

The mass spectrometry proteomics data of the PDX models have been deposited to the ProteomeXchange Consortium via the PRIDE partner repository with the dataset identifier PXD059245. The transcriptomic data of the PDX models have been deposited in the NCBI Sequence Read Archive under BioProject accession number PRJNA1346066.

## AUTHOR CONTRIBUTIONS

HK, EJ, NWW, SII, GJG, and RS contributed to study concept and design. HK, DMC, EJ, JWS, AuMA, DM, JLT, AmMA, NWW, BL, and EHO completed the acquisition of data. Analysis and interpretation of data was performed by HK, DMC, EJ, JWS, AuMA, DM, JLT, AmMA, NWW, BL, EHO CBC, MJB, SII, GJG and RLS. HK and RLS drafted the manuscript. EJ, CBC, MJB, SII, GJG, and RLS critically revised the manuscript concerning important intellectual content. HK, DMC, EJ, JWS, and DM performed statistical analysis. DMC, EJ, JWS, AuMA, DM, JLT, AmMA, NWW, BL, CBC, MJT, AP, SII, and GJG contributed administrative, technical, and/or material support. RLS supervised the study.

## ACKNOWLEDGEMENTS

This research was supported by the Mayo Clinic Department of Surgery, the Mayo Clinic Center for Cell Signaling in Gastroenterology, the Mayo Clinic DERIVE program, the Mayo Clinic Hepatobiliary SPORE (NCI/NIH P50 CA210964) to GJG and RLS, a supplement for liver cancer infrastructure (NCI/NIH 5P30 CA15083-43C1) to GJG and RLS, the NCI (1K08CA236874) to SII, the NIH (P30 DK084567), the NCI (U01CA271410 and P30CA15083) to AP, the Nijbakker-Morra Foundation to HK, and Fulbright Commission the Netherlands to HK. Schematic diagram figures were created with BioRender.com. KPPC cells were kindly gifted by Nabeel Bardeesy, PhD (Harvard Medical School, Massachusetts General Hospital, Broad Institute). NXP900 compounds were provided by Nuvectis Pharma, Inc., Fort Lee, NJ, USA.

