## Supplemental Tables for "Multiomics-driven discovery of predictive biomarkers and strategies to overcome resistance to SFK-YAP inhibition in cholangiocarcinoma"

**Supplemental Table 1.** Clinical characteristics of PDX models.

| PDX model | Patient sex | Patient age | Subtype | Molecular testing | Mutations | Amplifications | Other |
|---|---|---|---|---|---|---|---|
| 135 | Male | 34 | iCCA | Mayo Clinic Genetic Panel | TP53 P278A<br>BRAF V600E | KMT2C<br>AK5<br>PALB2<br>MDM2 | |
| 179 | Male | 66 | iCCA | FoundationOne | ATM L2077fs*<br>APC I1307K<br>BCOR M461fs*<br>BCOR S830fs* | FLT3<br>LYN<br>MCL1<br>MYC<br>NTRK1<br>BCL2L2 (eq)<br>MYST3 (eq)<br>NFKBIA<br>NKX2-1<br>ZNF703 | |
| 185 | Male | 66 | dCCA | WES research laboratory | PBRM1 splice donor<br>FANCC missense | | |
| 261 | Male | 65 | iCCA | WES research laboratory | LAMA2 missense<br>TPTE2 frameshift<br>ARID1A frameshift | | |
| 535 | Male | 78 | iCCA | Tempus | NRAS Q61<br>AR L110P<br>EGF R213Q<br>ETV4 V417fs<br>RNF43 G417E | | FGFR1 overexpression<br>FGFR2 overexpression<br>AR overexpression<br>MSS<br>TMB 2.6 |
| 194 | Male | 49 | pCCA | FoundationOne | TP53 E271*<br>CUL3 E533Q (subclonal)<br>STAT1 missense<br>FGFR3 missense<br>IDH1 missense<br>PBRM1 missense | ERBB2<br>FGFR3 (eq)<br>TOP2A (eq) | |
| 115 | Male | 58 | iCCA | Mayo Clinic Genetic Panel | IDH1 R132G<br>MET T1010I (VUS)<br>MSH2 missense<br>IDH1 missense<br>FOXP1 stop gain | | |
| 276 | Female | 58 | iCCA | FoundationOne | IDH2 R172T<br>PBRM1 2779+1G>A (splice site)<br>TP53 del(5-9) | | |
| 283 | Female | 54 | iCCA | FoundationOne | BAP1 D399fs*<br>FGFR2-AFF4 fusion | MCL1 | |
| 42 | Male | 36 | iCCA | WES research laboratory | FAT1 missense<br>SMAD4 missense<br>KRAS missense<br>ARID1A frameshift<br>PBRM1 splice donor<br>TP53 stop gain | | |
| 717 | Female | 45 | iCCA | Tempus | BRAF V600E<br>KEAP1 E205*<br>KRAS G12A<br>TP53 K320fs | | |

**Supplemental Table 2.** Primer sequences.

| Gene | Forward primer sequence (5' to 3') | Reverse primer sequence (5' to 3') |
|---|---|---|
| 18S | CGCTTCCTTACCTGGTTGAT | GAGCGACCAAAGGAACCATA |
| Human CTGF | GCAGCGGAGAGTCCTTCCAG | GGGCCAAACGTGTCTTCCAG |
| Human CYR61 | GAGTGGGTCTGTGACGAGGAT | GGTTGTATAGGATGCGAGGCT |
| Human NUAK2 | GATGCACATACGGAGGGAGATT | ATCACGATCTTGCTGCTGTTCT |
