## Supplemental Figures for "Multiomics-driven discovery of predictive biomarkers and strategies to overcome resistance to SFK-YAP inhibition in cholangiocarcinoma"

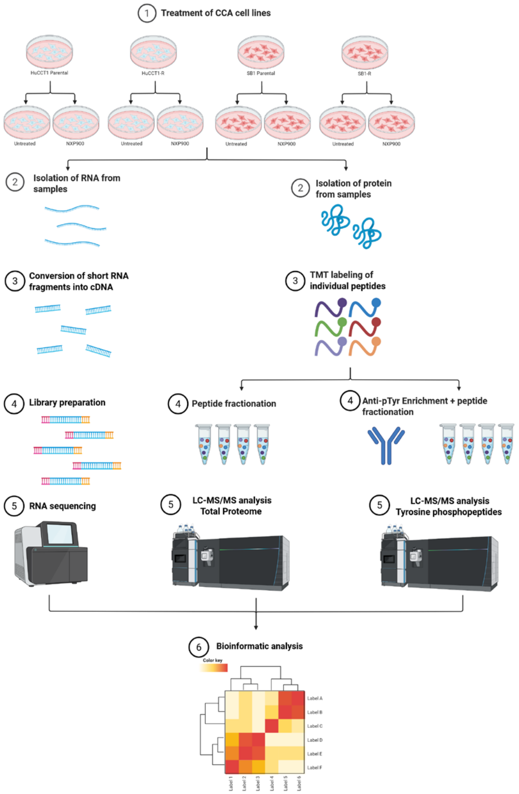


**Supplemental Figure 1.** Workflow schematic for transcriptomic, global proteomic and phosphoproteomic profiling in parental and resistant HuCCT1 and SB1 cells subjected to NXP900 treatment (1 μM for 6 hours) or vehicle.

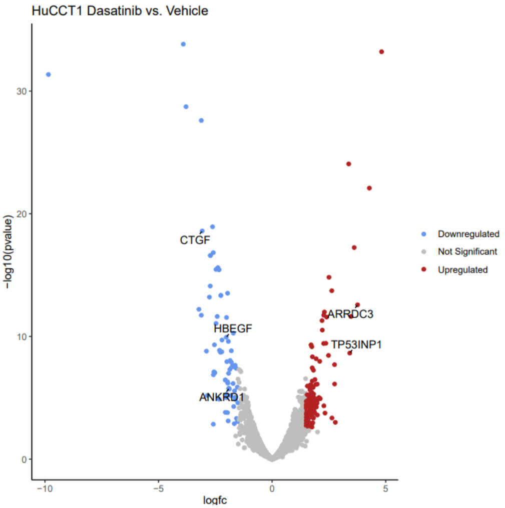


**Supplemental Figure 2.** Volcano plot comparing RNA-Seq gene expression (3 technical replicates/group) between HuCCT1 cells treated with dasatinib (1 μM for 6 hours) compared to vehicle. Significantly differentially expressed genes (–1.5 < logFC > 1.5, FDR < 0.05) highlighted blue (downregulated) and red (upregulated) with top genes of interest identified on plot.

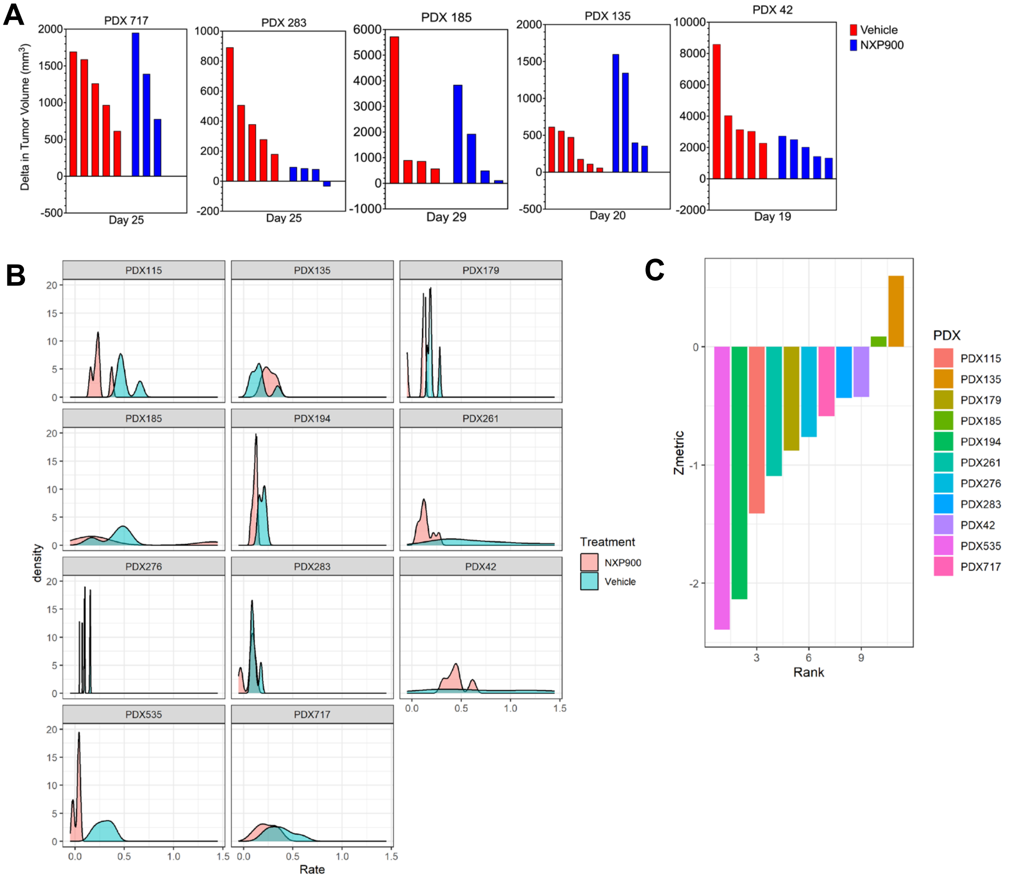


**Supplemental Figure 3.** (**A**) PDX tumor volumes at first day of significance or last day of treatment in non-responding models. (**B**) The per day growth rate (cm/day) for replicates of PDX models treated with NXP900 (red) or vehicle (blue) plotted by PDX model. (**C**) The summarized growth rate metric for each PDX model as a function of the difference in the growth rate distribution for NXP900 treated replicates and the growth rate distribution of vehicle replicates.

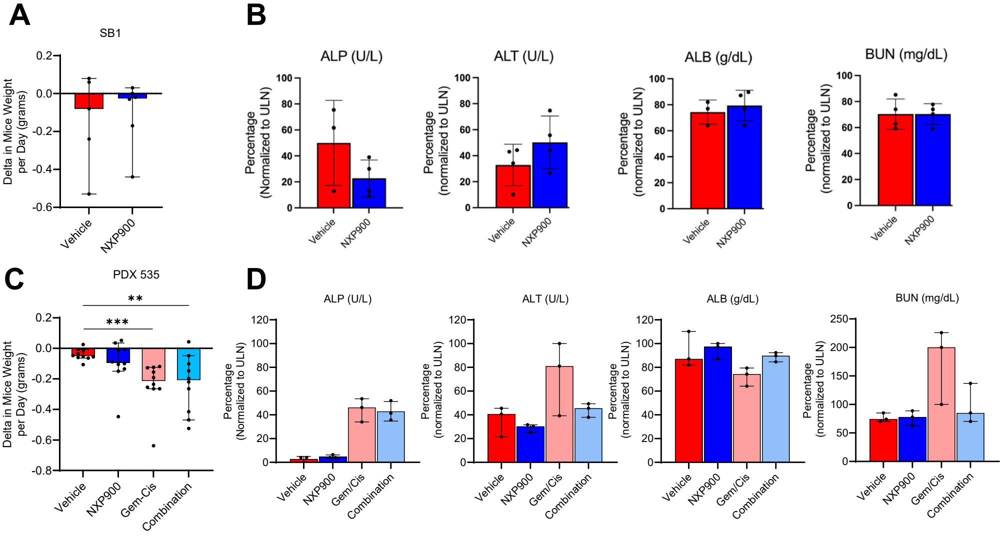


**Supplemental Figure 4.** (**A**) Delta in mice weight (g) per day in vehicle or NXP900-treated C57BL/6J mice with SB1 tumors following 2 weeks of treatment. (**B**) Serum liver and renal chemistries in vehicle or NXP900-treated C57BL/6J mice with SB1 tumors (n = 3). (**C**) Delta in mice weight (g) per day in vehicle, NXP900, gemcitabine-cisplatin, or combination-treated NOD/SCID mice with PDX 535 tumors. (**D**) Serum liver and renal chemistries in NOD/SCID mice with PDX 535 tumors treated with vehicle or NXP900 (n = 3). Data are shown as median with 95% CI. (**P < 0.01, ***P < 0.001). Statistical analysis was performed with 2-tailed Student t test.

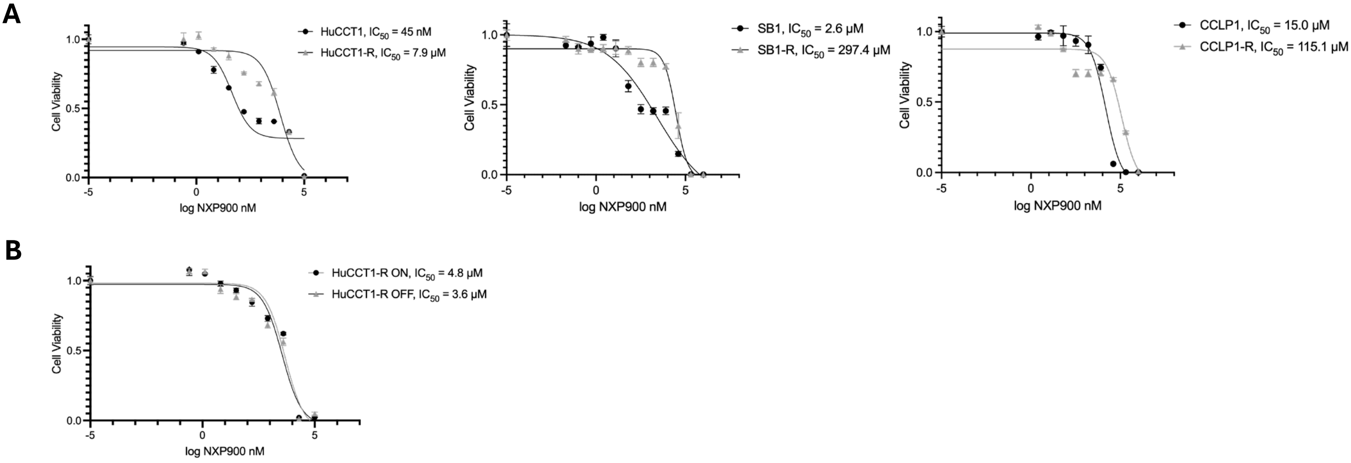


**Supplemental Figure 5.** (**A**) Viability dose-response curves and calculated IC50s of parental and resistant CCA cell lines treated with NXP900. IC50's were calculated at 24 hours (SB1) or at 48 hours of treatment (HuCCT1 and CCLP1). (**B)** Viability dose-response curves and calculated IC50s of the HuCCT1-resistant cell line cultured in the presence or absence (10 passages) of sublethal concentrations of NXP900. IC50's were calculated at 48 hours of treatment.

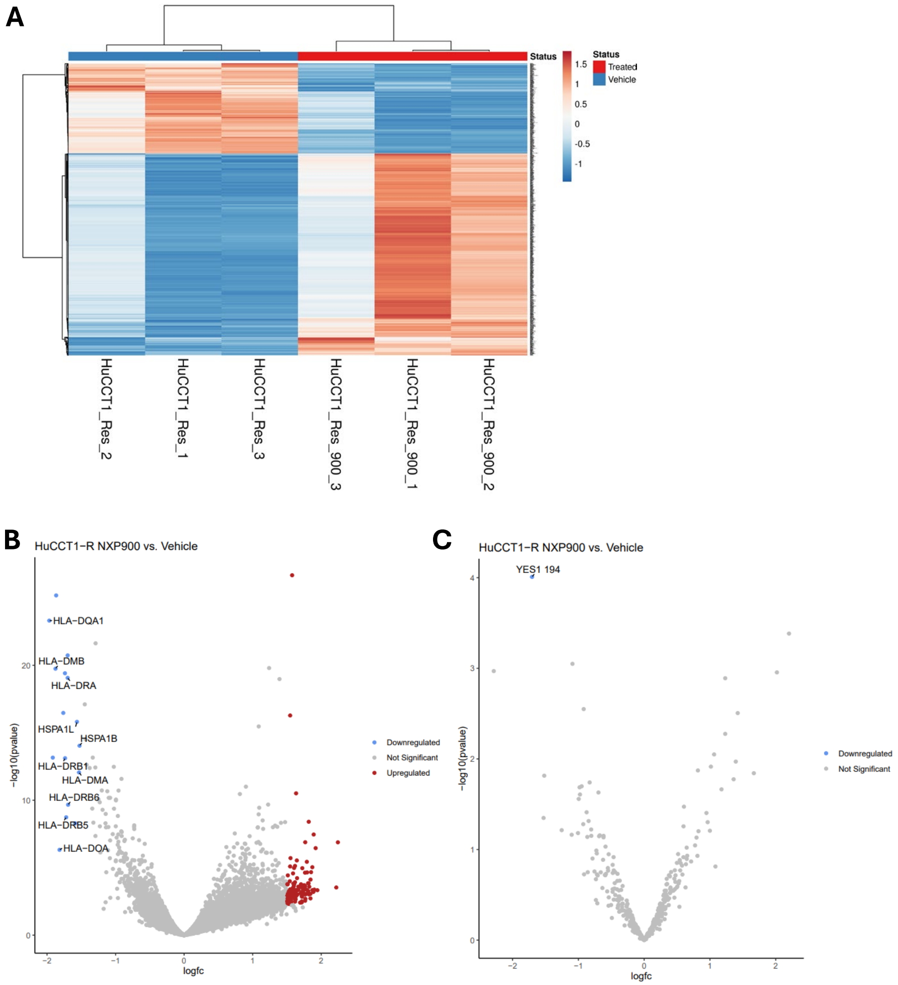


**Supplemental Figure 6.** (**A**) RNA-Seq of HuCCT1-resistant cell lines treated with NXP900 (1 µM for 6 hours) compared to vehicle. (**B**) Volcano plot comparing global proteomic abundance in vehicle- and NXP900-treated HuCCT1-resistant cells (3 technical replicates/group). Significantly differentially expressed phosphosites (–1.5 < logFC > 1.5, FDR < 0.05) highlighted blue (downregulated) and red (upregulated) with top proteins of interest identified on plot. (**C**) Volcano plot comparing phosphoproteomic (pTyr) abundance in vehicle- and NXP900-treated HuCCT1-resistant cells (3 technical replicates/group). Significantly differentially expressed phosphosites (–1.5 < logFC > 1.5, FDR < 0.05) are highlighted.

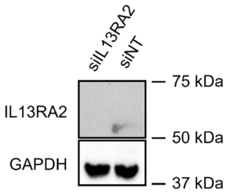


**Supplemental Figure 7.** Cell lysates from the HuCCT1-R cell line with siIL13RA2 or siNT for 48 hours subjected to immunoblot for IL13RA2. GAPDH was used as a loading control.
